# CDK4 Restricts Triple-Negative Breast Cancer Cell Migration via Phosphorylation-Driven Activation of Myo9b RhoGAP Function

**DOI:** 10.1101/2025.08.06.668850

**Authors:** Kanishka Parashar, Laia Simo Riudalbas, Arianna Ravera, Silvia Prieto Banos, David Moi, Barney F. Drake, Jialin Shi, Sarah Geller, Christophe Dessimoz, Georg E. Fantner, Dorian V. Ziegler, Lluis Fajas

## Abstract

Cyclin-Dependent Kinase 4 (CDK4) is a key regulator of cell cycle progression, driving the G0/G1-to-S phase transition through phosphorylation of Retinoblastoma 1 (RB1). Clinically, CDK4/6 inhibitors are under investigation in Triple Negative Breast Cancer (TNBC), a subtype characterized by invasiveness, aggressiveness and poor prognosis. While CDK4 is primarily targeted for its role in proliferation, emerging evidence suggests it may also regulate other cellular processes. In particular, the mechanisms by which CDK4 could influence cancer cell migration, remain largely unexplored, particularly in highly heterogenous cell line like MDA-MB-231. This study investigates whether CDK4 contributes to the regulation of TNBC cells migration and identifies the pathways involved in MDA-MB-231 cells, independently of its role in proliferation. We demonstrate that loss or inhibition of CDK4, using respectively CRISPR/Cas9 mediated CDK4 knockout and pharmacological CDK4/6 inhibitor, leads to enhanced migration capacities and reorganization of actin subcellular networks. Mechanistically, the absence of CDK4 results in decreased phosphorylation of Myo9b at serine 1935 (S1935), which enhances RhoA signaling, a key driver of cytoskeletal dynamics, leading to polarity defects and increased cell migration. These findings reveal a non-canonical function of CDK4 in limiting TNBC cell migration through the CDK4/CyclinD-Myo9b-RhoA signaling axis. This work highlights the broader cellular roles of CDK4 beyond its established function in proliferation and suggest that inhibition of Myo9b-RhoA pathway could reduce metastatic behaviour in TNBC treated with CDK4/6i, thereby informing future co-therapeutic strategies against aggressive cancer subtypes.

## Introduction

Cyclin-dependent kinases (CDKs) are highly conserved serine/threonine kinases that orchestrate cell cycle progression through phosphorylation of key substrates. Notably, CDK4, in complex with cyclin D drives the G0/G1-to-S phase transition by phosphorylating retinoblastoma protein (RB), releasing E2F transcription factors to promote DNA replication (1–3). Cyclin D types or CDK4 is often overexpressed in Triple Negative Breast Cancer (TNBC), driving uncontrolled proliferation, though CDK4/6 inhibitors reduce growth in RB1-proficient cells, with resistance common due to RB1 loss or bypass mechanisms (4). Dysregulation of the cyclin D-CDK4/6 axis is a hallmark of cancers, making CDK4/6 inhibitors (e.g., palbociclib) a cornerstone of therapy for patients with advanced estrogen receptor positive (ER)+/human epidermal growth factor receptor 2 (HER2)-negative breast cancer (5, 6). However, their efficacy in TNBC, a subtype lacking ER, progesterone receptor (PR), and HER2 expression, remains limited due to TNBC’s aggressive, heterogenous, and metastatic nature, which accounts for 10–15% of all breast cancer cases (7, 8). In addition to its well-established role in cell cycle regulation, CDK4 exerts non-canonical functions in different cancer models such as metabolism (9), autophagy (10), apoptosis (11), cellular senescence (12) and cytoskeletal dynamics ((1, 13). In recent years, several studies have further highlighted the role of the CDK4-cyclin D complex in facilitating key processes in cancer progression, such as migration (14), invasion (15) and metastasis (16), potentially through phosphorylation of cytoskeletal components or epithelial-to-mesenchymal transition (EMT) regulators (1, 17–20).In fact, CDK4 may influence cytoskeletal dynamics by modulating actin remodeling and microtubule stability, processes that are critical for cell migration, adhesion and intracellular transport (1, 13). However, the specific role of CDK4 in TNBC cell migration, particularly in highly metastatic cell lines like-MDA-MB-231, remains poorly understood. Key gaps include the molecular mechanisms by which CDK4 regulates migration, such as whether it phosphorylates migration-related proteins like Rho GTPases, and its interaction with cytoskeletal signaling pathways. Moreover, the effect of CDK4/6 inhibitors on TNBC migration, versus proliferation, is rarely studied, raising questions, if they could paradoxically enhance migration in resistant cells. These unanswered questions highlight the need to elucidate CDK4’role in TNBC migration and metastasis.

Cell migration, a hallmark of metastasis, is a complex process stimulated by receptor interactions (21, 22), which is highly controlled by members of the Ras homologous (Rho) family of small guanosine triphosphatases (GTPases), including Rho, Rac and Cdc42 (23–25). Rho GTPases regulate cytoskeletal dynamics by cycling between GDP-bound inactive and GTP-bound active states, controlling actin remodeling, actomyosin contractility, and the reorganization of cell-cell and cell-matrix adhesions during cellular migration (25, 26). As aberrant Rho GTPase signaling is a key driver of metastasis and cancer progression (27), modulating the expression of GTPase-activating proteins (GAPs), which are negative regulators of Rho GTPases, represents a promising yet underexplored strategy to reduce cancer cell migration and invasion. The interplay between CDK4 and GAPs in the context is largely unexplored, representing a critical research gap. Understanding whether CDK4 modulates Rho GTPase activity via phosphorylation of GAPs could reveal novel therapeutic targets to curb TNBC metastasis.

In this study, we aimed to elucidate CDK4’s role in TNBC cell migration using MDA-MB-231 cell line. Through CRISPR/Cas9-mediated CDK4 knockout, pharmacological CDK4/6 inhibition, and phospho-proteomics analysis, we investigated whether CDK4 regulates migration by modulating cytoskeletal dynamics in a phosphorylation-dependent manner. By addressing these gaps, this work seeks to uncover a novel CDK4-Myo9b-RhoA axis and assess its potential as a therapeutic target to limit TNBC metastasis.

## Results

### CDK4 inhibition or knockout increases cell size, induces topographic changes and promotes migration of TNBC cells

To study the role of CDK4 in TNBC cell proliferation, we first aimed to determine whether CDK4 inhibition or knockout affects cell morphology and migration by using MDA-MD-231 TNBC cells with CRISPR-Cas9-mediated CDK4 knockout (CDK4KO) (11). Compared with CDK4 wildtype (CDK4WT) cells, CDK4KO cells exhibited only slightly reduced proliferation (Sup. Fig. 1a), which is consistent with a distinct non-proliferative function of CDK4 in these cells. Using two independent clones, we next explored the effect of CDK4 on cytoskeleton-related phenotypes, focusing on cell size and morphology. Compared with CDK4WT cells, CDK4KO cells presented a significant increase in cell area and perimeter (Fig. 1a-b). This increase in cell size was confirmed over cell culture passages using a Trypan blue exclusion assay (Sup. Fig. 1b). Another independent CDK4KO clone (clone 18) exhibited a similar increase in cell size (Sup. Fig. 1c-d). Moreover, pharmacological inhibition of CDK4/6 with the CDK4/6 inhibitor palbociclib (Palbo) in CDK4WT cells partially recapitulated this phenotype, with a milder effect (Sup. Fig. 1e-f). These results indicated that CDK4 is required to maintain size of these cells.

**Figure 1.**
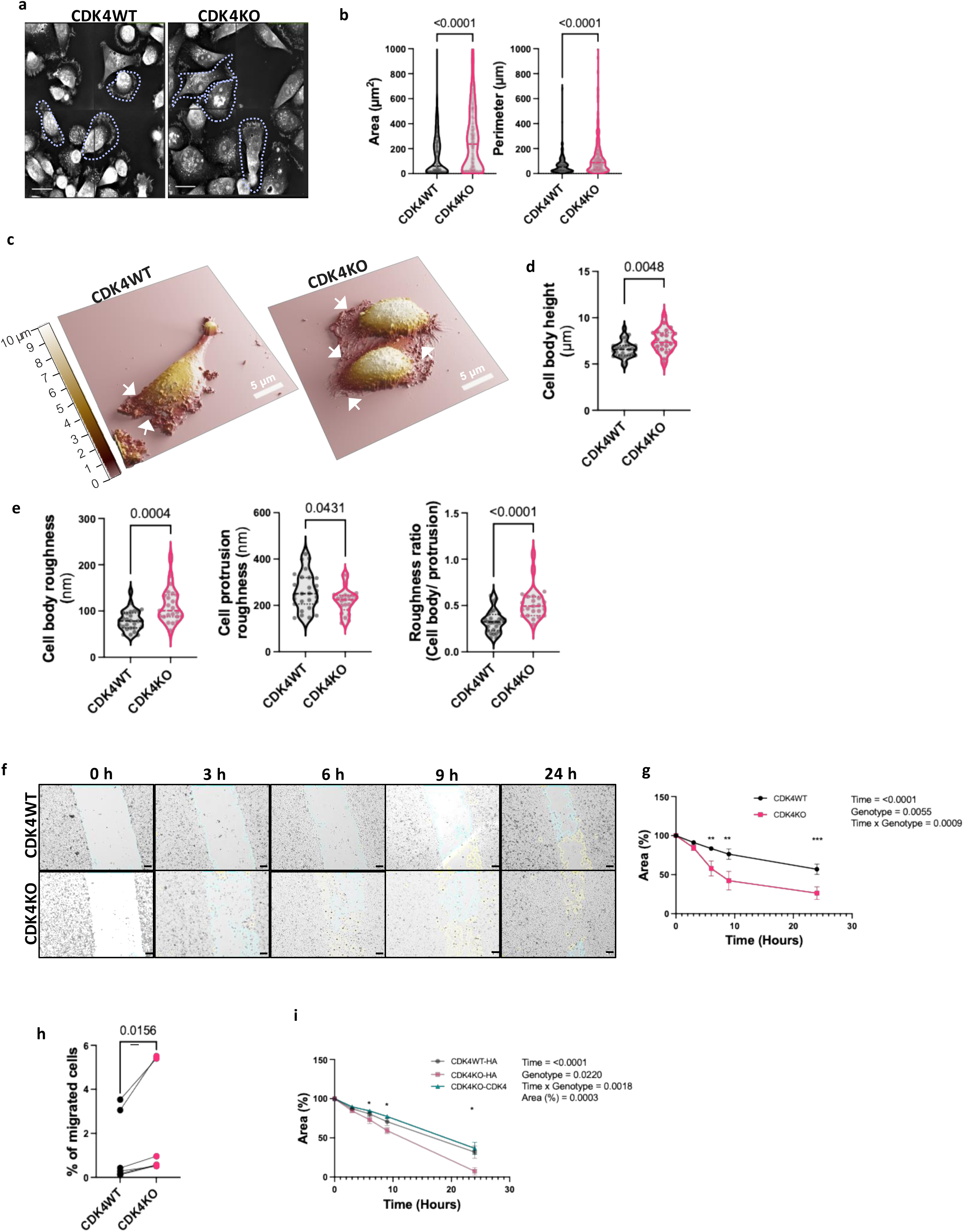
CDK4 inhibition or knockout (KO) increases cell size, induces topographic changes in and promotes migration TNBC cells. **a.** Representative image of CDK4WT and CDK4KO MDA-MB-231 TNBC cells from live movies. Scale bars: 30 μm, with purple lines indicating cell boundaries **b.** Quantification of Eve software (From Nanolive microscope) analysis for cell area and perimeter in CDK4WT and CDK4KO cells. N= 4 independent biological replicates from n= 313 (WT) and 335 (KO) cells. Unpaired T-test, Mann-Whitney test. **c.** Representative SICM 3D images illustrating cell topography from CDK4WT and CDK4KO MDA-MB-231 cells. Color scale: cell height from 0 to 10 μm. **d-e**. Measurement of cell height (d), roughness (e) of CDK4WT and CDK4KO cells over cell body cell protrusion, with ratio between the two on a cell-by-cell basis. **f.** Representative pictures (Sharp contrast) of CDK4WT and CDK4KO MDA-MB-231 TNBC cells from a scratch assay at 0, 3, 6, 9, and 24 hours after wounding, scale bar: 100 μm. **g.** Quantification of the scratch assay results as a percentage of the area covered by CDK4WT and CDK4KO cells with respect to time in hours. Mixed effects analysis, Tukey’s multiple comparison test. The data are shown as the mean +/− SEM of N= 3 independent biological replicates from n= 9 biological replicates. **h**. Quantification of the percentage of CDK4WT and CDK4KO MDA-MB-231 TNBC cells that migrated through Transwell membranes with a pore size of 8 μm. Paired t test, N= 6 independent biological replicates. **i.** Quantification of the migration of CDK4WT and CDK4KO MDA-MB-231 TNBC cells transfected with an empty plasmid containing HA (CDK4WT-HA & CDK4KO-HA, respectively) and CDK4KO TNBC cells transfected with a CDK4-expressing plasmid (CDK4KO-CDK4) by the scratch assay. 2-way ANOVA, Tukey’s multiple comparison test. The data are shown as the mean +/− SEM of N= 3 independent replicates from n= 4 biological replicates. Exact p values are displayed.

In order to study further the cell structure, we next examined the cell topology of the cells. Scanning ion conductance microscopy (SICM) (28), which measures cell height and roughness revealed the presence of membrane protrusions projected upward in the z axis in CDK4 KO TNBC cells (arrows, Fig. 1c-d), suggesting the formation of ruffle-like structures in these cells. Further topological analysis indicated that CDK4KO cells were taller and had a rougher surface over the cell body but smoother surface than wild type cells over peripheral cell extensions (Fig. 1e).

Changes in cytoskeleton organization is associated with the migration ability of the cells. We therefore assessed cell migration over 24 hours using scratch and transwell assays in our models. Interestingly, the percentage of scratched area progressively decreased over time, with a significant faster closure in CDK4KO cells compared to CDK4WT cells, indicating enhanced migratory capacity CDK4KO cells (Fig. 1f-g-)), transwell migration assay confirmed a significant increase in the percentage of migrated CDK4KO compared to control (Fig. 1h). The independent CDK4KO clone 18 and CDK4WT cells treated with the CDK4/6i palbociclib covered the scratched area sooner than their respective control conditions (Sup. Fig. 1g-h), indicating a similar enhanced migratory cell phenotype. Most importantly, partial rescue of CDK4 expression in CDK4KO cells, using an exogenous CDK4 plasmid, was sufficient to limit cell migration (Fig. 1i and Sup. Fig. 1i), restoring the WT phenotype. Collectively, these findings suggest that CDK4 is a key regulator of TNBC cells motility, with its absence/inhibition leading to enhanced migratory behaviour.

### CDK4KO disrupts the directional migration and front-rear polarity of actin localization in TNBC cells

To better understand the migration pattern of the cells, we utilized the Nanolive microscope to create live cell migration movies, which were used to track the cell migration. The Cartesian trajectory plot revealed distinct patterns of migration of CDK4WT and CDK4KO cells (Fig. 2a and Sup. Fig. 2a). Indeed, the movement of CDK4WT cells was rather uniform, suggesting stable migration behavior, whereas the trajectories of CDK4KO cells showed greater variability in the pattern and direction of migration, indicating altered cell migration dynamics. The trajectory deviation plot (Fig. 2b and Sup. Fig. 2a) further highlighted these differences, revealing that the deviation area was narrow for CDK4WT cells, indicating predictable movement, but broader and more irregular for CDK4KO cells, reflecting increased random motion and instability in cell migration patterns. Furthermore, time-lapse imaging of CDK4KO cells provided further insight into the nature of this random migration, revealing a distinct, contractility-driven migration cycle (Fig. 2c, Sup. Video 1-2). Protrusions initially extended from the cells, forming contractile tails, and the tails then underwent rapid displacement, repeating this cycle (Fig. 2c, Sup. Video1-2). Similar random contractile migration patterns were observed for the independent CDK4KO clone (clone 18) and Palbociclib-treated CDK4WT cells (Sup. Fig. 2b-c). These observations suggest that the absence of CDK4 significantly impacts cell migration patterns, potentially through changes in cytoskeletal organization or motility regulation.

**Figure 2.**
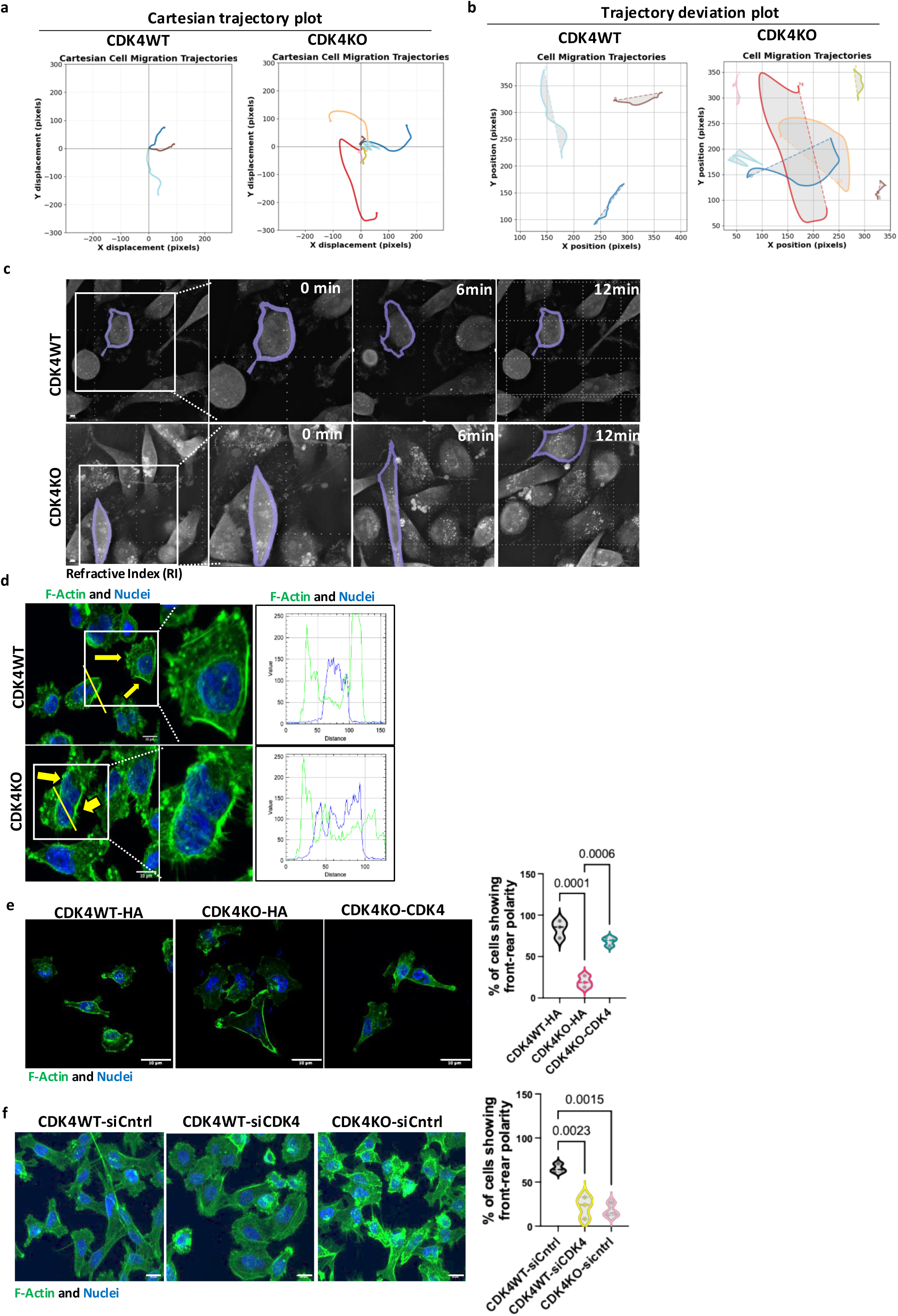
CDK4KO disrupts the directional migration and cell front-rear polarity of actin localization in TNBC cells. **a**. CDK4WT and CDK4KO MDA-MB-231 cell’s cartesian trajectories across the experimental conditions, showing the representative paths of multiple cells. Each line corresponds to the path of a single cell, and the lines are aligned at the origin to allow comparison of migration patterns. **b**. Trajectory deviation from a straight migration path. The original cell trajectory (solid line) is plotted alongside the straight-line displacement from the initial to the final position (dashed line). The area between these two paths (in light gray) quantifies the deviation from directional movement, and the corresponding value is reported**. c.** Snapshots (Sharp contrast) of migrating CDK4WT and CDK4KO MDA-MB-231 TNBC cells from movies showing the time (in min), with the analyzed cells in purple. The white square represents the cropped cells, and the dotted lines show the movement of the cells across time. Scale bar: 30 μm. **d.** Representative images (Sharp contrast) of CDK4WT and CDK4KO MDA-MB-231 TNBC cells showing small F-actin protrusions (green) and nuclei (blue), Scale bar: 10 μm. **e.** Representative images (Sharp contrast) of CDK4WT and CDK4KO TNBC cells transfected with an empty plasmid expressing HA (CDK4WT-HA & CDK4KO-HA, respectively) and CDK4KO TNBC cells transfected with a CDK4-expressing plasmid (CDK4KO-CDK4), Scale bar: 10 μm, with associated quantification. One-way ANOVA, Tukey’s multiple comparison test, N= 3 independent biological replicates. **f.** Representative images (Sharp contrast) of CDK4WT and CDK4KO TNBC cells transfected with siCntrl (CDK4WT-siCntrl & CDK4KO-siCntrl, respectively) and CDK4WT TNBC cells transfected with siCDK4 (CDK4WT-siCDK4) Scale bar: 10 μm, with associated quantification. One-way ANOVA, Tukey’s multiple comparison test. The data are shown by violin plot, showing all points and each point represents one biological replicate, N= 3 independent replicates. Exact p values are displayed.

To determine whether CDK4 influences the migration phenotype of TNBC cells through changes in cytoskeletal organization or motility regulation, we analyzed cell morphology with a focus on F-actin filament distribution, a key determinant of cell migration, as this parameter modulates cell shape and migration (29–31) by fixed cell imaging where F-actin was stained by phalloidin (green). CDK4WT cells exhibited pronounced front-rear polarity, characterized by dense actin localization at the leading edge (Fig. 2d). In sharp contrast images, CDK4KO cells displayed a loss of front-rear polarity and uniform distribution of actin across the cell membrane (yellow arrow, Fig. 2d). Most importantly, the disruption in actin organization was reversed by the restoration of CDK4 expression in CDK4KO cells (Fig. 2e). Furthermore, this actin reorganization was also observed siRNA-mediated CDK4 silencing in CDK4WT cells (Fig. 2f and Sup. Fig. 2d) and independent CDK4KO clone (clone 18) (Sup. Fig. 2e). Altogether, these findings suggest that CDK4 is critical for maintaining actin front-rear polarity and that the loss of CDK4 contributes to altered migration patterns.

### CDK4 inactivation increases the expression of actin-associated proteins, altering actin dynamics and promoting filopodium formation in TNBC cells

To explore the molecular basis of altered cell size and migration upon the loss of CDK4, we utilized previously published proteomics data in CDK4WT and CDK4KO MDA-MB-231, TNBC cells (32). We notably observed a change in the expression of proteins involved in focal adhesion, cell migration, the actin cytoskeleton and organization (Fig. 3a-b). The intensity of protein expression, represented on the y-axis, shows a central clustering of highly protein expression (Fig. 3a). These results indicate a substantial impact of CDK4 deletion on the proteomic profile of MDA-MB-231 cells. Subsequent analysis of protein expression changes showed the differential expression of cytoskeleton-associated proteins in CDK4KO versus WT MDA-MB-231 cells. These findings suggest that CDK4 deletion not only alters global protein expression but also impacts cytoskeleton-related functions, potentially influencing cellular structure and motility in MDMA-MB-231 cells (Fig. 3b).

**Figure 3.**
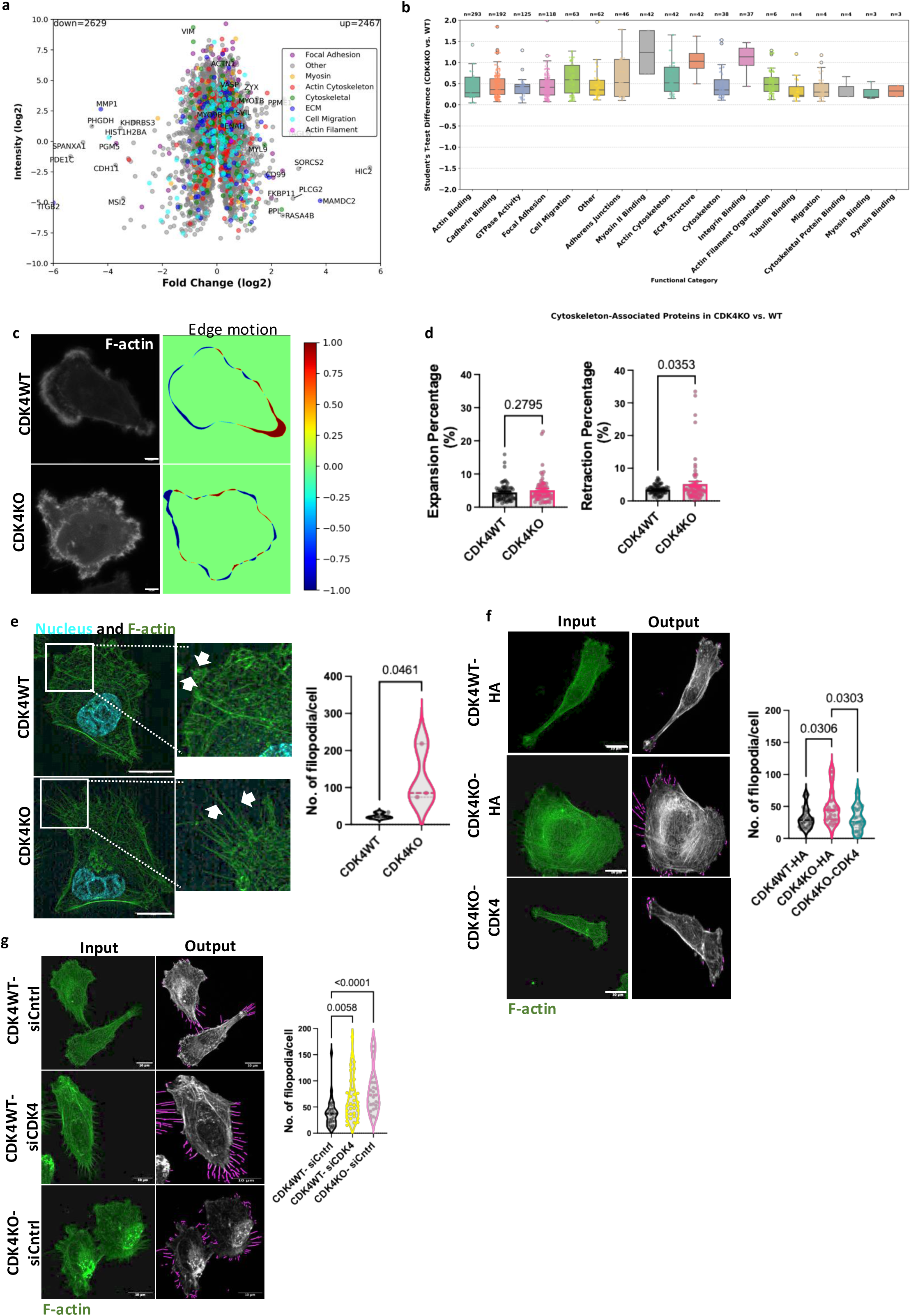
CDK4 inactivation increases actin-associated proteins levels, altering actin dynamics and promoting filopodium formation in TNBC cells. **a.** Volcano plot depicting log2 fold change versus log10 intensity, highlights the relative abundance of proteins that are down- and upregulated from the proteomics data from CDK4WT and CDK4KO MDA-MB-231 cells (32) **b.** Bar plot of the difference in the enrichment of the differentially expressed proteins between the CDK4KO and CDK4WT groups in functional categories according to Student’s t test. **c.** Edge motion and actin density maps. Positive values in the edge motion map (in red) indicate regions of frequent expansion, whereas negative values (in blue) indicate areas of frequent retraction over time. The actin density map shows regions of consistent actin enrichment, with a higher intensity corresponding to areas where high actin concentrations were repeatedly observed throughout the sequence, Scale bar: 10 μm. **d.** Quantification of edge dynamics across experimental groups. Bar plots show the average percentage of cell area undergoing expansion and retraction for each condition, enabling comparison of protrusive and contractile behavior between genotypes. Unpaired Student Tt test. N= 3 independent biological replicates, n= 30-40 cells approx. **e.** Representative images (sharp contrast) of CDK4WT and CDK4KO MDA-MB-231 TNBC cells showing small F-actin extensions (filopodia, green), nuclei (cyan). Associated quantification the number of filopodia per cell in the images; each dot represents one cell. Unpaired T-test. **f**. Representative images (Sharp contrast) and associated quantification output showing filopodia in pink lines of CDK4WT and CDK4KO TNBC cells transfected with an empty plasmid expressing HA (CDK4WT-HA & CDK4KO-HA, respectively) and CDK4KO TNBC cells transfected with a CDK4-expressing plasmid (CDK4KO-CDK4). Quantification of the number of filopodia per cell from images. Kruskal-Wallis test, Dunn’s multiple comparisons test. N= 3 independent biological replicates; each dot represents one cell, approx. n= 30-40 cells. **g.** Representative images (Sharp contrast) and associated quantification output showing filopodia in pink lines of CDK4WT and CDK4KO TNBC cells transfected with siCntrl (CDK4WT-siCntrl & CDK4KO-siCntrl, respectively) and CDK4WT TNBC cells transfected with siCDK4 (CDK4WT-siCDK4). Quantification of the number of filopodia per cell. Kruskal-Wallis test, Dunn’s multiple comparisons test. N= 3 independent biological replicates, n= 30-40 cells, approx. Exact p values are displayed.

Based on the results of the proteomics analysis. we next investigated the dynamics of the actin cytoskeleton, which is a critical regulator of cell motility (33). We stably expressed an F-actin reporter plasmid in CDK4WT and CDK4KO MDA-MB-231 cells to monitor actin dynamics at the cell periphery. Edge motion was analyzed by quantifying extension and retraction movements across four quadrants of the cell membrane. Motion vectors and subsequent inward and outward movements were analyzed (arrows, Sup. Fig. 3a, Sup. Video 3-4). Compared with CDK4WT cells, CDK4KO cells presented increased actin dynamics, characterized by an increased number of motion vectors with a greater average distance and angular dispersion at each quadrant (Sup. Fig. 3b). Specifically, membrane retraction was significantly increased in CDK4KO cells compared to CDK4WT cells, whereas changes in membrane extension were not significantly different (Fig. 3c-d).

During F-actin staining in fixed and live cell imaging, we also observed increased filopodia in CDK4KO cells, consistent with their role in cell migration and altered focal adhesion dynamics (34) (35). After quantifying the number of extensions observed (arrows) per cell, we found that CDK4KO cells presented a significantly greater number of filopodia than CDK4WT cells (Fig. 3e). This phenotype was reversed by the reintroduction of CDK4 into CDK4KO cells (pink lines, Fig. 3f and Sup. Fig. 3c) and was recapitulated by silencing CDK4 in CDK4WT cells (pink lines, Fig. 3g and Sup. Fig. 3d). Moreover, in another CDK4KO clone (clone 18) (Sup. Fig. 3e) and CDK4WT cells treated with CDK4/6 inhibitor palbociclib (Sup. Fig. 3f), also showed an increased number of filopodia per cell compared to their respective controls. In addition to an increased number of filipodia, CDK4KO cells displayed longer filopodia (Sup. Fig. 3g) in CDK4-silenced WT cells and CDK4/6i-treated CDK4WT cells (Sup. Fig. 3h). Collectively, these results demonstrate that CDK4 plays a pivotal role in maintaining front-rear polarity and regulating filipodium formation by modulating actin-associated proteins, thereby enhancing the migratory behaviour of TNBC cells.

### CDK4KO enhances RhoA activity and ROCK-dependent migration, with Myo9b phosphorylation as a CDK4 target

Based on CDK4 function as a kinase, we hypothesized that it may modulate cell migration by phosphorylating proteins involved in the regulation of the actin cytoskeleton and migration-related processes. Interestingly, enrichment analysis of downregulated genes in CDK4KO MDA-MB-231 cells form phosphoproteomic data (32) by Metascape (36) revealed significant enrichment in the Rho GTPase cycle signaling pathway (Fig. 4a) suggesting that the absence of CDK4 impairs Rho GTPase-mediated processes, potentially affecting cytoskeletal dynamics and cell migration. Leading from this, RhoA is known to regulate actin polymerization, contractility and adhesion via ROCK1/2 and phosphorylated myosin light chain (pMLC) (37, 38)) (Fig. 4b). To assess RhoA activity, we performed directly a colorimetric assay and quantified indirectly pMLC levels by immunoblotting. CDK4KO cells presented significantly elevated RhoA activity (Fig. 4c) and increased expression of pMLC, a downstream marker of RhoA activity (Fig. 4d). These changes were consistent with those observed in another CDK4KO clone (clone 18) (Sup. Fig. 4a). Taken together, these results clearly indicate that CDK4 loss enhances RhoA activity.

**Figure 4.**
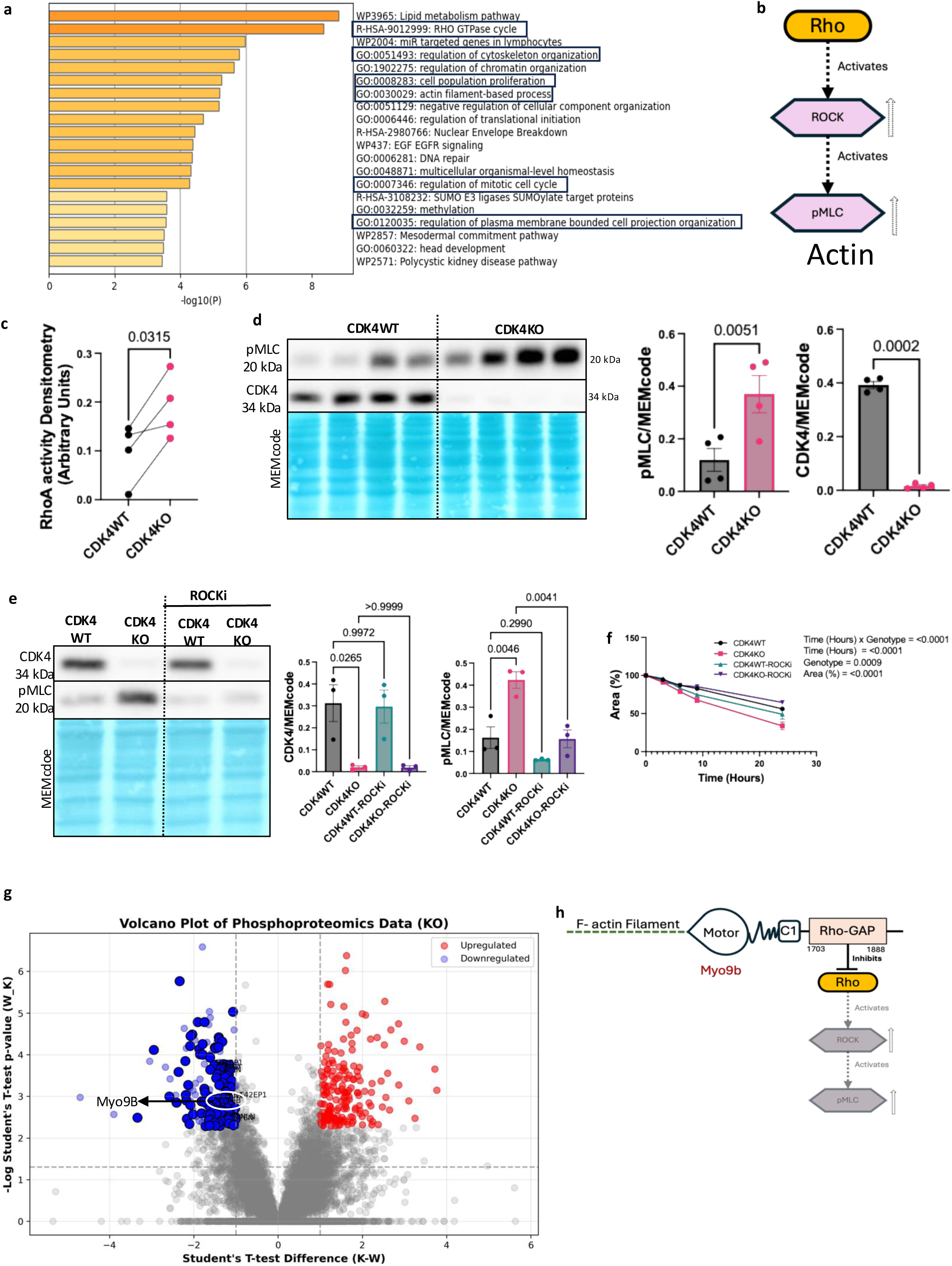
CDK4KO enhances RhoA activity and ROCK-dependent migration, with Myo9b phosphorylation as a CDK4 target. **a.** Enrichment analysis of phosphoproteomic data generated by Metascape (36) highlighting the enrichment of the differentially expressed proteins in the Rho GTPase cycle, cytoskeleton organization and the actin filament process. **b.** Schematic showing RhoA and downstream targets**. c.** Densitometric analysis in arbitrary units of RhoA activity in CDK4WT and CDK4KO MDA-MB-231 TNBC cells. Paired t test. N= 4 independent biological replicates. **d.** Immunoblots and relative protein levels of RhoA, pMLC, CDK4, and MEMcode in CDK4WT and CDK4KO TNBC cells. Unpaired t test. Data are shown as the mean +/− SEMs of N = 4 independent biological replicates. **e.** Immunoblots and relative protein levels of pMLC, CDK4 and MEMcode in CDK4WT and CDK4KO TNBC cells non-treated and treated with ROCK inhibitor (20 μM) for 24 hours. One-way ANOVA, Tukey’s multiple comparison test. Data are shown as the mean +/− SEMs of N = 3 independent biological replicates. **f.** Quantification of scratch assay as a percentage of the area covered analysis for CDK4WT and CDK4KO TNBC cells non-treated and treated with ROCK inhibitor (20 μM) for 24 hours. 2way ANOVA, Tukey’s multiple comparison test. Data are shown as the mean +/− SEMs of N = 3 independent biological replicates and n=6 biological replicates. **g.** Volcano plot of phosphoproteomic data showing log-transformed Student’s t test p values vs. Student’s t test differences, showing upregulated (red) and downregulated (blue) proteins in CDK4KO cells vs. CDK4WT cells. **h.** Schematic of Myo9b and a potential downstream pathway. Exact p values are displayed.

To address whether ROCK1/2 pathway was involved in the phosphorylation of pMLC and subsequent enhanced migration of CDK4KO cells, we treated these cells with a classical ROCK inhibitor Y-27632 (20 μM) for 24 hours. This treatment notably reduced pMLC expression in CDK4KO cells to level comparable to those observed in CDK4WT cells, as well as migration rates (Fig. 4e-f), indicating that ROCK-mediated signaling pathway functionally contributes to the migration of cells lacking CDK4.

To investigate how CDK4 is responsible for the change in RhoA signaling observed upon deletion, we further analyzed phosphoproteomic data (32) to identify both putative CDK4-specific phosphorylation targets and regulators of RhoA activity (Fig. 4g). We also used the kinase prediction tool from PhosphoSitePlus (39) to determine the percentile scores and ranks of specific sites, reflecting the likelihood of their phosphorylation by CDK4 (Sup. Fig. 4b, Sup. Table S4). Among the identified putative CDK4 targets was Myo9b, a Rho GTPase-activating protein (Rho-GAP) known to inhibit RhoA activity via its Rho-GAP domain and subsequently regulate cell shape, contractility and migration (40–45) (Fig. 4h). Phosphoproteomic analysis revealed reduced phosphorylation of Myo9b at serine 1935 (S1935), a site bearing the CDK4-recognized SPXK/R/P motif, in CDK4KO cells (Fig. 4g and raw data (32)). Also, the kinase percentile score for S1935 was notably high (96.498) (Sup. Fig. 4b-c), indicating a strong likelihood of CDK4 involvement because of SP motif. Additionally, a nearby serine at position 1926, which also fits the potential CDK4 phosphorylation motif, was considered for kinase score calculation, although not appeared in phosphoproteomics data. However, the kinase prediction percentile score for S1926 was lower (84.885) (Sup. Fig. 4c), suggesting weaker CDK4 association. Apparently, both S1926 and S1935 are conserved across species (Sup. Fig. 4d), highlighting their evolutionary significance. These findings suggest that CDK4-mediated phosphorylation of Myo9b at S1935 may allosterically enhances Myo9b RhoGAP activity, thereby suppressing RhoA signaling.

### Phosphorylated Myo9b at S1935 regulates RhoA activity *via* the interaction between Myo9b-RhoGAP and RhoA

Myo9b modulates RhoA activity through direct interaction of its RhoGAP domain with RhoA (46). To investigate this potential impact of phosphorylation at S1926 and S1935 on Myo9bRhoGAP-RhoA interaction, we employed AlphaFold3 (47). The predicted structures of Myo9b (Sup. Fig. 5a) and the Myo9b-RhoA complex (Fig. 5a) displayed relatively high proportions of disordered regions (0.35 and 0.34, respectively, Sup. Fig. 5b). In contrast phosphorylation at either S1926, S1935 or at both sites resulted in more ordered structures, with disordered region proportions of 0.04, 0.02, and 0.03 (Fig. 5b-c). Notably, only in the Myo9b-RhoAcomplex structure containing Myo9b phosphorylated at S1935 there was an interaction between the RhoGAP domain of Myo9b and RhoA (Fig. 5d), with interacting residues at 4.5 to 6 Å apart (Sup. Table S1). However, the domains are not in contact with no phosphorylation (>14 A°), S1926 phosphorylation (>20 A°), or dual S1926/S1935 phosphorylation (>20 A° Fig. 5e, Sup. TableS2). Furthermore, proximity ligation assays (PLAs) revealed and confirmed that the number of interactions between Myo9b and RhoA was decreased in CDK4KO cells compared to CDK4WT cells (Fig. 5f and Sup Fig. 5c). Altogether, these results suggest that CDK4-mediated phosphorylation of Myo9b at S1935 stabilizes the Myo9b-RhoA multimer structure, allowing binding to RhoA and the modulation of RhoA activity.

**Figure 5.**
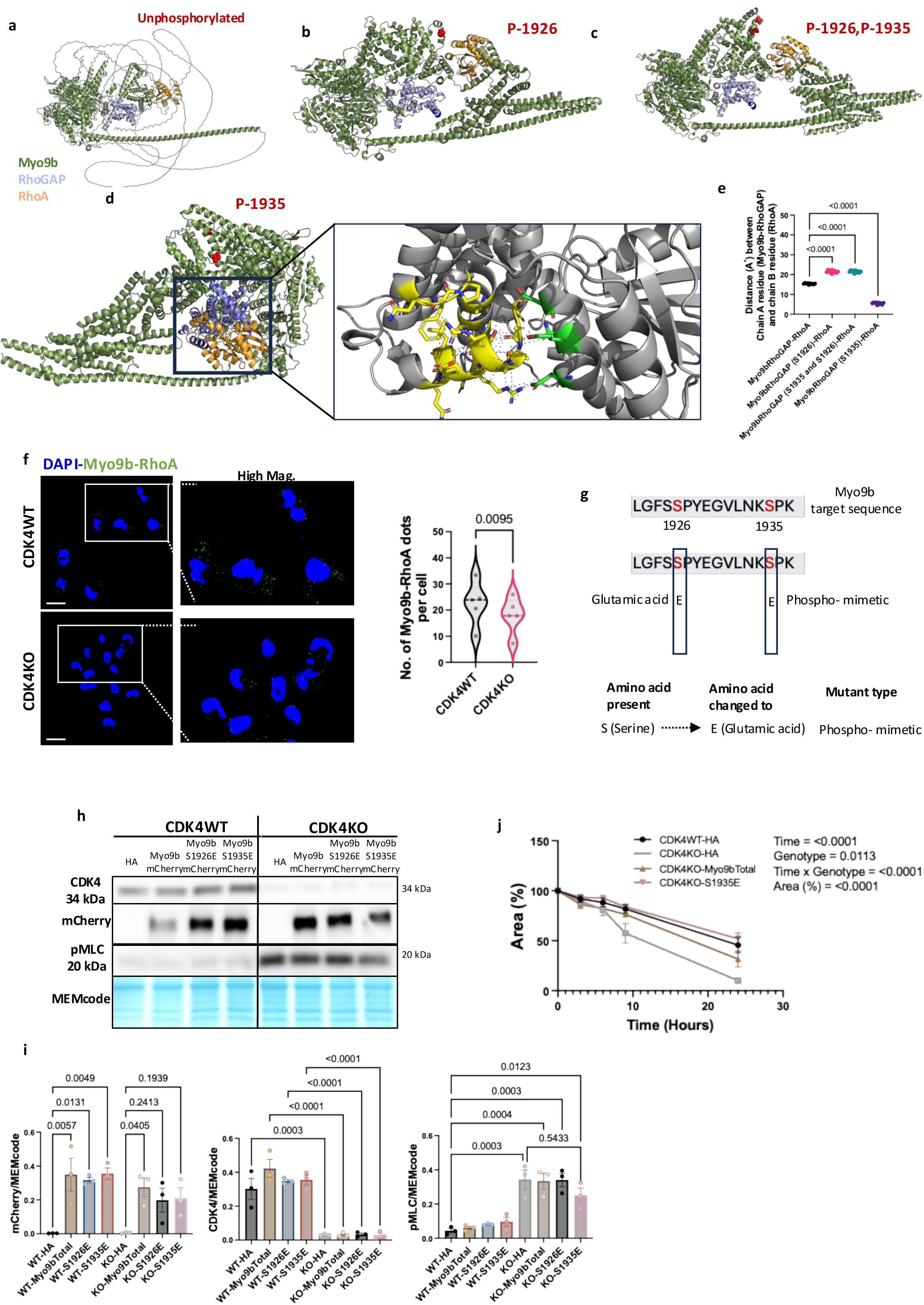
Phosphorylated Myo9b at S1935 regulates RhoA activity *via* the interaction between Myo9b-RhoGAP and RhoA. **a.** Structural prediction of human Myo9b and the RhoA multimer by AlphaFold. Myo9b is shown in green and purple (RhoGAP domain), and RhoA is shown in orange. **b-c.** Structures of the phosphorylated forms of the Myo9b-RhoA complex (top left: P-1926; top right: P-1926 and P-1935; bottom: P-1935). **d.** Zoomed-in view of the RhoGAP interaction site and RhoA. The interaction site and interacting atoms (<= 4 Å) are shown in stick representation in addition to the ribbon diagrams and colored by element (blue=N, red=O). The carbon atoms of RhoGAP are shown in yellow, and those of RhoA are shown in green. The dashed lines represent polar contacts between the RhoGAP interaction site and any atom**. e.** Distance between the 10 closet contacts of Myo9b’s RhoGAP domain with RhoA residues for the four multimer versions (Sup. TableS2). One-way ANOVA, Tukey’s multiple comparison test. **f.** Proximity ligation assay (PLA) using Myo9b and RhoA antibodies in CDK4WT and CDK4KO TNBC cells. Representative images and associated quantification of Myo9-RhoA dots per cell are shown. Scale bars: 10 μm.N = 5 independent biological replicates, n= approx. 60-80 cells. Two-sided paired t test. **g** Schematic representation of the Myo9b peptide sequence and a phosphomimetic mutant. **h-i.** Immunoblots and relative protein levels of mCherry, CDK4, pMLC and MEMcode in CDK4WT and CDK4KO MDA-MB-231 TNBC cells transfected with empty plasmid (HA), the Myo9b mCherry plasmid, the Myo9b S1926E mCherry plasmid and the Myo9b S1935E mCherry plasmid for both CDK4WT and CDK4KO MDA-MB-231 cells. 2way ANOVA, Tukey’s multiple comparison test. Data are shown as the means +/− SEMs of N = 3 independent biological replicates. **j.** Quantification of the scratch assay results as a percentage of the area covered by CDK4WT and CDK4KO cells transfected with HA empty plasmid and Myo9b total and Myo9b-S1935E mutated plasmid, with respect to time in hours. 2way ANOVA, Tukey’s multiple comparison test. The data are shown as the mean +/− SEM of N= 3 independent biological replicates from n= 5 biological replicates. Exact p values are displayed.

To confirm that CDK4 regulates Rho-ROCK-pMLC pathway involved in cell motility through Myo9b phosphorylation, Myo9b expression was silenced in CDK4WT cells using siRNA under basal condition, followed by the introduction of phosphomimetic form of Myo9b. CDK4WT cells silenced for Myo9b exhibited increased pMLC levels shown qualitatively (Sup. Fig. 5d), disrupted front-rear polarity (Sup. Fig. 5e) and displayed a random migration pattern (Sup. Fig. 5f) compared to control conditions, similar to previous observations in CDK4KO cells. Together, these data underscore the role of Myo9b in maintaining polarized migration in MDA-MB-231 cells. Moreover, reintroduction of the phosphomimetic mutant Myo9b S1935E, but not Myo9b S1926E, into CDK4KO cells (Fig. 5g), confirmed by mCherry tag detection, led to a partial reduction of pMLC levels compared to control CDK4KO cells (Fig. 5h-i). These findings indicate that CDK4-mediated phosphorylation of Myo9b at S1935 enhances Myo9b RhoGAP activity, suppressing RhoA activity. Accordingly, and most importantly, the migration capacity of CDK4KO cells decreased after reintroducing the phosphomimetic mutant Myo9b-S1935E plasmid by transfection (Fig. 5j), suggesting the importance of this phosphosite in the inhibition of migration.

Collectively, these data demonstrate that CDK4-mediated phosphorylation of Myo9b at S1935 enhances the interaction between the Myo9b-RhoGAP domain with RhoA, thereby suppressing RhoA activity and limiting TNBC cells migration. In contrast, reduced Myo9b phosphorylation at S1935 in the absence of CDK4, disrupts this regulatory mechanism, leading to increased RhoA activity and enhanced cell migration.

## Discussion

This study, investigates the role for CDK4 in Triple Negative Breast Cancer (TNBC) cell migration, focusing on its non-canonical functions.

Our findings uncover a previously unexplored role for CDK4 in TNBC cell migration, distinct from its classical function RB-dependent cell cycle progression. In MDA-MB-231 cells, CDK4 deletion or pharmacological inhibition with palbociclib reduced proliferation but enhanced migratory capacity, accompanied by significant cytoskeletal remodeling, including loss of cell front-rear polarity and increased filopodium formation. These findings align with emerging evidence of the involvement of CDK4’s involvement in cytoskeletal dynamics (1, 13, 48, 49) and support the “go or grow” hypothesis, which posits an inverse relationship between cancer cell proliferation and migration (50, 51). This dichotomy, observed in TNBC and other cancers, suggest that reduced proliferative activity may enhance migratory potential, a critical step in metastasis (52–55). Notably, enhanced migration following CDK4 deletion and CDK4/6 inhibition mirrors observation in lung and pancreatic cancers (56, 57), though context-specific effects exist either promote or suppress breast cancer migration and metastasis, highlighting the need to clarify CDK4’s role in TNBC’s heterogenous subtypes (58). In contrast to previous reports, however, a study also suggest that CDK4/6 inhibition or cyclin D1 expression increases breast cancer migration (59), whereas other studies indicate that CDK4/6 inhibition or cyclin D1 expression may suppress metastasis (18, 19). These discrepancies underscore the complex, context-dependent role of CDK4 pointing to the influence of cancer cell type and the hormonal context of CDK4/6 inhibition on cancer cell migration, showing a critical gap in its understanding the molecular mechanism and therapeutic implications. However, our findings contribute to addressing this gap by demonstrating that CDK4 suppresses migration in MDA-MB-231 cells.

Moreover, CDK4KO cells exhibited disrupted directional migration, potentially reflecting increased cell motility driven, for example, through alterations in the extracellular matrix (ECM) or cellular energy status (60). The enhancement of filopodium formation in CDK4KO cells further supports increased migration, which is consistent with studies linking induced filopodium formation to enhanced migration in MDA-MB-231 cells (61, 62). Additionally, espin-driven filopodium formation in confined spaces promotes metastasis of TNBC cells (35).

Our study identified a new target of CDK4, Myo9b, which modulates downstream RhoA activity. Myo9b, a unique actin-based motor protein with a RhoGAP domain, inactivates RhoA by converting it from its GTP-bound to GDP-bound state, regulating actin cytoskeleton dynamics critical for cell migration (41, 63–66). Myo9b influences the shape and motility of various cell types, including macrophages, immune cells, and Caco2 (BBE) intestinal epithelial cell monolayers (41, 42, 64, 67–71). However, the structural basis for the role of Myo9b in regulating RhoA activity remains unclear. In CDK4WT MDA-MB-231 cells, the phosphorylation of Myo9b at S1935 enhanced Myo9b RhoGAP activity, suppressing RhoA activity to maintain front-rear polarized actin organization and directional migration. In contrast, in CDK4KO cells, the absence of Myo9b S1935 phosphorylation leads to RhoA hyperactivation, elevated ROCK1/2 signaling as shown in PamGene kinase activity profiling (32), and subsequently increased pMLC levels, promoting contractility and increasing migration. Furthermore, RhoA activation has also been shown to increase filamentous actin and myosin II contractility (72). This CDK4-Myo9b-RhoA axis addresses a critical gap in understanding how CDK4 integrates the cytoskeletal signaling pathways, such as Rho GTPase signaling, to regulate TNBC migration. This finding also informs the gap regarding CDK4’s non-cell cycle roles, revealing a novel phosphorylation target that links cell cycle and migration pathways. Besides, RhoA’s bidirectional interplay with cell cycle regulators, such as cyclin D1, suggests a complex cross-regulation between proliferation and migration. In 3T3 fibroblasts, RhoA was found to be essential for G1 progression via cyclin D1; however, the dynamic regulation of RhoA activity across the cell cycle has not been fully explored (73, 74). RhoA also plays critical roles in mitosis and cytokinesis, particularly at the cleavage furrow, where it drives actomyosin ring contraction and abscission (75, 76). RhoA activity fluctuates during the cell cycle, peaking during G2/M to facilitate mitotic processes, as evidenced by its regulation by GEFs such as Ect2 and GAPs such as p190RhoGAP (77, 78). Remarkably, RhoA and its effectors (ROCK2, myosin II, LIMK, and cofilin) influence G1-S phase progression via p21 and cyclin D1 expression (79), linking to a bidirectional relationship between RhoA and cell cycle regulators. Our study thus contributes to better understand the cross-regulation of cell cycle with cytoskeleton dynamics. Moreover, our data improves our knowledge about the migration of MDA-MB-231TNBC cells, involving CDK4 in this process.

The identification of Myo9b S1935 for the first time, as a putative CDK4 phosphorylation site highlights its role in modulating RhoA activity, with structural remodeling confirming enhanced Myo9b-RhoA interaction upon phosphorylation at S1935. Interestingly, reintroduction of S1935 phosphomimetic plasmid in CDK4KO cells reduced the migration rate, emphasizing the critical role of this site in cell migration. While PhosphoSitePlus has predicted multiple Myo9b phosphorylation sites, including S1354 - whose phosphorylation enhances Rho-GTP suppression in human platelets (40) - the relevance of S1935 is uniquely highlighted in our study.

While our study provides valuable insights into CDK4’s role in TNBC migration, several opportunities for further exploration remain. We have yet to fully characterize the compensatory mechanisms of RhoA activity or the functional role of other Myo9b phosphorylation sites, which could offer additional insights into cytoskeletal dynamics. Future validation in diverse TNBC models, including patient-derived xenograft, would strengthen the generalizability of our findings. Moreover, the context-dependent effects of CDK4/6 inhibitors across TNBC subtypes require further investigation to guide targeted therapeutic strategies, contributing to a deeper understanding of subtype-specific and *in vivo* effects. Exploring these avenues could further illuminate CDK4’s multifaceted role in TNBC progression and enhance its therapeutic potential.

Our study establishes CDK4 as a pivotal regulator of TNBC cell migration, extending its role beyond cell cycle regulation and offering new insights into its non-canonical functions. Given the widespread use of CDK4/6 inhibitors in breast cancer treatment, our findings underscore the potential risk of enhanced tumor dissemination in TNBC, an aggressive subtype lacking targeted therapies due to absence of ER, PR, and HER2 expression. While chemotherapy, remains the primary treatment for TNBC, its efficacy is often constrained by resistance and tumor heterogeneity, highlighting the urgent need for novel strategies. The identification of the CDK4-Myo9b-RhoA axis presents a promising strategy to suppress TNBC migration while controlling proliferation, addressing critical gaps in understanding CDK4’s role in metastasis. Indeed, CDK4/6 inhibitors and ROCK inhibitors have been studied individually in terms of their effects on cancer cell proliferation and migration, respectively, no studies have explored their combined impact on TNBC. Identification of the CDK4-Myo9b-RhoA/ROCK axis in TNBC migration regulation here suggests that such combination therapies targeting both proliferation (CDK4/6 inhibitors) and migration (RhoA/ROCK/Myo9b specific inhibitor), could potentially overcome the limitations of current treatments. Elucidating these synergistic effects may open new avenues for therapeutic development in TNBC.

Future research should explore how CDK4 interacts with other cytoskeletal regulators across diverse cancer types and evaluate Myo9b phosphorylation as a potential therapeutic target to reduce TNBC’s metastatic potential. These efforts could pave the way for innovative, precision-based therapies, transforming the management of this challenging disease.

## Materials and methods

### Cell culture and reagents

Triple-Negative Breast Cancer (TNBC) MDA-MB-231 cells (American Type Culture Collection ATCC, HTB-26™) were cultured in Roswell Park Memorial Institute (RPMI)-1640 medium with GlutaMAX (Gibco™, Ref: 61870036) supplemented with 10% heat-inactivated FBS (HyClone, Cat. No: SV30160, Lot: RB35956), 10 mM HEPES (Gibco™, Ref: 15630122) and 1 mM sodium pyruvate (Gibco™, Ref: 11360070). Cells were maintained at 37°C in a humidified atmosphere with 5% CO2. All the cells were routinely tested and confirmed mycoplasma-free using standard PCR-based assays. CDK4KO MDA-MB-231 were generated using CRISPR/Cas9 gene editing as previously described (11). Pharmacological treatments were performed using Palbociclib (Merck, Sigma-Aldrich, Ref: PZ0383-5MG-PD0332991) and ROCK inhibitor Y-27632 (Abcam, Ref: ab120129).

### Plasmid transfection and antibodies

Empty-HA (Ctrl) and CDK4-WT plasmids were kindly provided by Dr. Marcos Malumbres (CNIO, Spain). The mCherry-Myo9b plasmid was a gift from Dr. Martin Bähler (Addgene plasmid #134918; http://n2t.net/addgene:134918; RRID:Addgene_134918, (42)), and the pEGFP-C1 Lifeact-EGFP was a gift from Dr. Dyche Mullins (Addgene plasmid #58470; http://n2t.net/addgene:58470; RRID:Addgene_58470, (80)). Cells were seeded one day prior to transfection. Transfection was carried out using X-tremeGENE™ HP DNA at a 2:1 (μL:μg) ratio, following a 20-minute incubation in Opti-MEM (Gibco™, Ref: 51985034) before addition to the cells. Stable cell lines expressing Lifeact-EGFP for visualization of F-actin in (green) were generated by antibiotic (600 µg/mL G418) for 2 weeks post transfection.

### Scratch assay

MDA-MB-231, CDK4WT and CDK4KO cells were cultured in 6 well plates (Corning Incorporated Costar^®^ (Ref: 3516) until reaching 95–100% confluency. A linear scratch was gently introduced across the cell monolayer, using a sterile micropipette tip, the wells were subsequently rinsed with PBS 1X to remove cellular debris. For pharmacological inhibition assay, the cells were pre-treated for 24 hours with Palbociclib (1 μM) and the ROCK inhibitor-Y-27632 (20 μM) before a scratch was induced. Brightfield (BF) images were acquired over a 24-hours period using the CELENA X high-content imaging system (Logos Biosystems, CX30004) with 4x objective. Wound closure was quantified using Fiji (Fiji-ImageJ2, Version:2.14.0/1.54p) with Wound_healing_size_tool pluging (81) and results were expressed as the percentage of the wound area over time.

### Transwell migration assay

MDA-MB-231 cells (25,000-50,000/well) were seeded into the upper chamber of 8-μm pore Transwell insert plate (Corning, Costar, Ref: 3422) in 150 μl of serum free medium. The lower chamber contained 600 μl of medium with (10% FBS) as chemoattractant. After 18-20 hours incubation at 37°C and 5% CO2, non-migratory cells were removed from the upper membrane surface with a cotton swab. Migrated cells were fixed with 4% paraformaldehyde (Thermo Fisher, Ref: 28906) for 10 minutes at Room Temperature (RT) and stained with Hoechst 33342 (Thermo Fisher, Ref: Hoechst 33342) for 10 min at RT, and followed by two washes with PBS1X. Images were acquired using the CELENA High-Content Imaging System (Logos Biosystems) with 4x objective in the DAPI channel, and, quantified by using Fiji (Fiji-ImageJ2, Version:2.14.0/1.54p) software.

### RhoA activity assay

MDA-MB-231, CDK4WT and CDK4KO cells were plated and allowed to grow for 24 hours, and then serum starved for 6 hours. RhoA-GTP levels were measured at RT using the RhoA G-LISA assay kit (Cytoskeleton Inc., Cat. # BK124-S) following the manufacture’s instruction. Briefly, active GTP-bound RhoA in cell or tissue lysates binds to immobilized Rho GTP-binding protein in a 96-well plate, while inactive GDP-bound Rho is washed away. Bound active RhoA is detected by a specific antibody and the activation levels are quantified by absorbance at 490 nm using a microplate spectrophotometer. Wells containing lysis buffer only were used as assay blank.

### Site-directed mutagenesis

The Myo9b-mCherry plasmid (Addgene plasmid #134918) was used to generate the phosphomimetic mutant (Myo9b-mCherry S1936E and/or/S1935E) using theQuick-Change Lightning Site-Directed Mutagenesis Kit (Agilent, Cat. # 210518), according to the manufacturer’s instructions.

### Immunoblotting

MDA-MB-231 cells were lysed in M-PER™ Mammalian Protein Extraction Reagent (Thermo Scientific, Ref: 78501) supplemented with 1X protease and phosphatase inhibitor cocktails (Thermo Scientific, Ref: 78429 and Ref: 1861277, respectively). Protein concentration was determined by Pierce™ Bradford assay (Thermo Scientific, Ref: 23200). Equal amounts of protein (15-20 μg) were separated on 6 or 12% SDS-PAGE and transferred onto nitrocellulose membranes (Amersham Protran, Ref: 10600002). The membranes were blocked with 1X TBS containing 0.05% Tween/5% dry non-fat milk powder for 1 h and incubated at 4°C with primary antibodies overnight. The next day, the membranes were washed 3 times with 1X TBS containing 0.05% Tween and further incubated with a secondary antibody diluted in 1X TBS containing 0.05% Tween/5% skim milk for 1 hour at RT. The membranes were then washed 3 times with 1X TBS containing 0.05% Tween, and the protein bands were detected using an enhanced chemiluminescence (ECL) kit (Amersham, Ref: RPN2235 and Advansta, Ref: K-12045-D20) and imaged by using FUSION FX7 imaging system (Vilber Lourmat, France). Antibodies details and dilutions are listed in supplementary table (Table S3. List of antibodies)

### Immunofluorescence labelling and microscopy

MDA-MB-231 cells were rinsed with 1X PBS and fixed with 4% methanol-free paraformaldehyde (Thermo Fisher, Ref: 28906). Cells were permeabilized with 0.3% Triton X-100 and then blocked for 1 hours at RT in blocking buffer (1X PBS+ 2% bovine serum albumin (BSA)+0.05% Tween 20)). The primary antibodies were diluted as indicated (Sup. TableS3.) in blocking buffer overnight at 4°C. After three washes times in blocking buffer for 5 min each at RT. The cells were incubated for 30 mints at RT in the dark with Alexa Fluor 488-conjugated donkey anti-mouse or donkey anti-rabbit (Jackson ImmunoResearch, respectively, Ref: 715-546-150; Ref: 711-546-152; and with Alexa Fluor 594-conjugated donkey anti-mouse or donkey anti-rabbit (Jackson ImmunoResearch, respectively Ref: 715-586-150; Ref: 711-586-152) diluted 1/500 together with Hoechst 33342 (1/10 000, In vitrogen Cat.No.H3570) in 1% BSA, followed by three 1X PBS washes. Sample were mounted with Fluoromount-G^®^ (SouthernBiotech: Cat. No. 0100-01) and stored at 4°C. Imaging was performed on Zeiss LSM 880 Airyscan confocal microscope and a Zeiss Elyra 7 SR-SIM microscope (1) equipped with a 63x oil immersion objective (Fig. 3e).

### Time-lapse microscopy

Label-free snapshot images and cell migration videos shown in Fig. 2c and Sup. Video 1 and 2, were acquired by using the 3D Cell Explorer Fluo microscope (Nanolive SA, Tolochenaz, Switzerland) with EVE Explore software.Cell morphology (area and perimeter) was analyzed using Nanoive’s EVE Analytics software. For images in Fig.3c and (Sup Videos 3 and 4), CDK4WT and CDK4KO cell lines stably expressing Lifeact-EGFP actin-green were cultured on poly-D-lysine coated MatTek glass bottom dishes (35-mm, 14-mm diameter) (MatTeK Corporation, Ref: P35gc-o-14-C). Imaging was performed on ZEISS LSM 880 microscope with Airy-scan detection using a 63x oil immersion objective, capturing frames every 5 seonds for 1 minute.

### Proximity Ligation Assay (PLA)

PLA was performed using the, Duolink^®^ reagent (Merck, Ref: DUO92014, DUO92002, DUO92004). Cells were washed with 1X PBS and fixed for 10 min in 10% paraformaldehyde. An equal volume of 1 M glycine was added, followed by a 15-min wash with 100mM glycine. Cells were permeabilized with 0.1% Triton-X-100 and incubated overnight at 4 °C with primary antibodies (Sup. Table S3). After washing with 1X PBS containing 0.3% Tween-20, cells were incubated with PLA probes. Ligation and polymerization steps were performed according to the manufacturer’s instructions (Merck, Duolink^®^ procedure). PLA puncta were quantified using the “Analyze particles” tool in Fiji software (Fiji-ImageJ2, Version:2.14.0/1.54p).

### Scanning ion conductance microscopy (SICM) imaging, data processing and analysis

Images from 4% PFA fixed MDA-MB-231 CDK4WT and CDK4KO cells were taken in Phosphate Buffered Saline (PBS), at RT, in ibidi imaging dish (Cat.No: 81156, μ-Dish 35 mm, high). Imaging was performed in hopping mode with bias voltages of 200mv with typical ion currents of 4nA and typical pipette tip inner diameters of 100nm. A detailed description of the homebuilt microscope platform has been published here (28). SICM images were processed using Gwyddion (82). Image defects with the maximum width of 1 pixel were removed by filling-in the neighboring line. The cells were marked by implementing height thresholding, with the height data of the images leveled by shifting the mean height value of the substrate part to zero. To analyze the roughness and height of the cell, we separated the small structure on the cell membrane surface from the cell topography by using frequency split or two polynomial degrees leveling. The cell bodies were masked by shrinking the mask of the cells. The root mean square roughness of a cell body and boarder were calculated from the high frequency image, and cell height was extracted by using the max value of the low frequency image. All SICM topography data were normalized by 13 µm and exported in 16-bit Portable Network Graphics format with gray and customized color bar. We used the open-source 3D rendering tool Blender 3D to generate the three-dimensional images of the cells. The corresponding SICM height data and colored data were imported and used as a heigh map and projected color on the topography.

### Image analysis

#### Trajectory extraction and visualization

To analyze cell migration dynamics, the position of each cell was tracked across time-lapse frames by computing its geometric center. This yielded a raw trajectory for each cell, represented as a sequence of time-stamped spatial coordinates. To ensure data quality, trajectories shorter than a defined minimum fraction of the experimental duration were discarded. The remaining trajectories were temporally smoothed to reduce noise and measurement variability. Smoothing was followed by the exclusion of abrupt displacements exceeding a defined threshold, which were likely due to segmentation errors or tracking mismatches. Each valid trajectory was then used to generate two types of plots. In the first representation, called the trajectory deviation plot (Fig. 2b), both the observed path and the idealized straight-line displacement (from the initial to final position) are displayed. The area enclosed between the two paths was computed and normalized by trajectory length to quantify migration persistence, with larger enclosed areas indicating more erratic movement. In the second representation, called the Cartesian trajectory plot (Fig. 2c), trajectories were translated to a common origin by aligning all the starting positions at (0,0). This Cartesian transformation allowed for the comparison of directionality across cells.

#### Edge motion and actin density

To quantify membrane and cytoskeletal dynamics over time, two complementary analyses were performed: one based on edge motion and the other on actin intensity. For edge motion analysis, binary cell masks from each time point were compared sequentially to identify temporal changes at the cell boundary. By subtracting the mask of each frame from that of the preceding frame, pixels that were either gained or lost at the periphery were detected. This allowed us to distinguish between expanding regions (pixels present only in the current frame) and retracting regions (pixels absent from the current frame but present in the previous frame). These frame-to-frame differences were accumulated over the entire time series to generate a spatial heatmap (Fig. 2c – Edge Motion) showing the frequency and location of protrusive and contractile events. To enable quantitative comparisons of membrane behavior across conditions, the number of expanding and retracting pixels was calculated for each cell and expressed as a percentage of the total cell area. These values were then used to generate a comparative plot (Fig. 2d) summarizing the balance of protrusion and retraction across the experimental groups, revealing genotype-dependent differences in edge dynamics. In parallel, to evaluate the spatial distribution of actin over time, an actin density map was constructed from the original fluorescence images. For each frame, high-intensity actin regions were isolated by applying a dynamic threshold based on the mean and standard deviation of nonzero pixel intensities. Only pixels with an intensity above this threshold were retained, highlighting areas of increased actin concentration. These threshold regions were then summed across all frames and normalized to the number of timepoints, producing a final map (Fig. 2d – Actin Density) that reflects regions of sustained or recurrent actin enrichment throughout the sequence.

#### Filopodia analysis

Filopodia number and length were measured using the semi-automated Filo Quant plugin (Fiji-ImageJ2, Version:2.14.0/1.54p),as shown in Sup. Fig. 3c-d.

### Computational analysis of the Myo9b and the Myo9b-RhoA multimer structures

We assessed changes in the structural conformations of Myo9b and the Myo9b-RhoA complex using the AlphaFold3 server (47). The amino acid sequences input into AlphaFold were obtained from UniProt with the IDs Q13459 (Myo9b) and P61586 (RhoA). First, we predicted the structure of Myo9b only, and then we separately predicted the structures of 1) Myo9b and the RhoA multimer, 2) Myo9b phosphorylated at S1926 and the RhoA multimer, 3) Myo9b phosphorylated at S1935 and RhoA, and 4) Myo9b phosphorylated at both S1926 and S1935 and RhoA. Phosphorylation was included as a posttranslational modification. The proportions of disordered regions for the four Myo9b-RhoA multimer versions were obtained from the output of AlphaFold3, and we predicted the disordered regions of Myo9b using AIUPred (83). The distances between residues in the predicted structures were calculated using the PDB module of Biopython (v. 1.85). Figures showing the structures were created with PyMol.

### Data representation and statistical analysis

Results are presented as,bar plots, showing individual values are presented as the means ± SEMs of N independent biological replicates for in vitro experiments. In whisker and violin plots, n indicates the number of cells. The Shapiro-Wilk normality test was applied to raw data before any other analyses. Other statistical analyses and tests that were performed are indicated in the figure legends. For comparisons between two groups of nonnormally distributed data, the nonparametric Mann-Whitney test was performed. For comparisons between two groups of normally distributed data, the following two-tailed parametric tests were performed: Student’s t test (equal variance) and Welch’s t test (nonequal variance). For comparisons among more than two groups, repeated measures (RM) (for paired data) or ordinary (for unpaired data) one-way analysis of variance (ANOVA) was used, and Tukey’s multiple comparisons test was subsequently performed. All the statistical analyses were performed using GraphPad Prism 9.1.0 software (ns: not significant; * p < 0.05; ** p < 0.01; *** p < 0.001; **** p < 0.0001).

## Data availability

PhosphoProteomics (https://proteomecentral.proteomexchange.org/cgi/GetDataset?ID=PXD046326. And Proteomics (https://proteomecentral.proteomexchange.org/cgi/GetDataset?ID=PXD046353) respectively (32). Any remaining data will be available from the corresponding author if needed.

## Acknowledgements

The authors would like to thank the lab members of Prof. Lluis Fajas Coll group for the helpful discussions, Manfredo Quadroni and the Proteomic Analysis Facility (PAF), Fabienne Lammers and the Genomic Technologies Facility (GTF), Arnaud Paradis and Luigi Bozzo at the Cellular Imaging Facility (CIF) Genopode and Agora respectively, and the Scientific Service at the Center for Integrative Genomics (SSC). This work was supported by the University of Lausanne (UNIL). We acknowledge the Swiss National Science Foundation (SNF) grants #31003A_143369 and #310030_207688 to L.F. group. SPV, DM, and CD group were supported by SNF grants #205085 and #216623. G.E.F. and B.F.D acknowledge the funding from the European Research Council under grant No. ERC-2017-CoG InCell, the EPFL Center for Imaging under grant No. 563292, and ETH domain - ETH Open Research Data (ETH-ORD) under grant No. 563386.

## Author contributions

K.P., D.V.Z. and L.F. conceived and designed the study. K.P. performed and analyzed experiments. A.R. performed the live movies image analysis. K.P., L.S.R., B.F.D, J.S, G.F. suggested, performed and analyzed the SCIM microscope data for cell topological study. K.P., S.P.B., D.M. and C.D. suggested, performed and analyzed the protein folding data from AlphaFold. K.P. and D.V.Z. performed and analyzed PLA experiments. K.P., D.V.Z. and L.F. wrote the manuscript. All authors approved the final version of the article.

## Declaration of generative AI and LLM

During the preparation of this manuscript the author(s) used ChatGPT, Grok in order to enhance clarity. After using this tool, the author(s) reviewed and edited the content as needed and take(s) full responsibility for the content of the published article.

## Supplementary information

Correspondence and requests for materials should be addressed to Lluis Fajas.

