## supplemental figures for "CDK4 Restricts Triple-Negative Breast Cancer Cell Migration via Phosphorylation-Driven Activation of Myo9b RhoGAP Function"

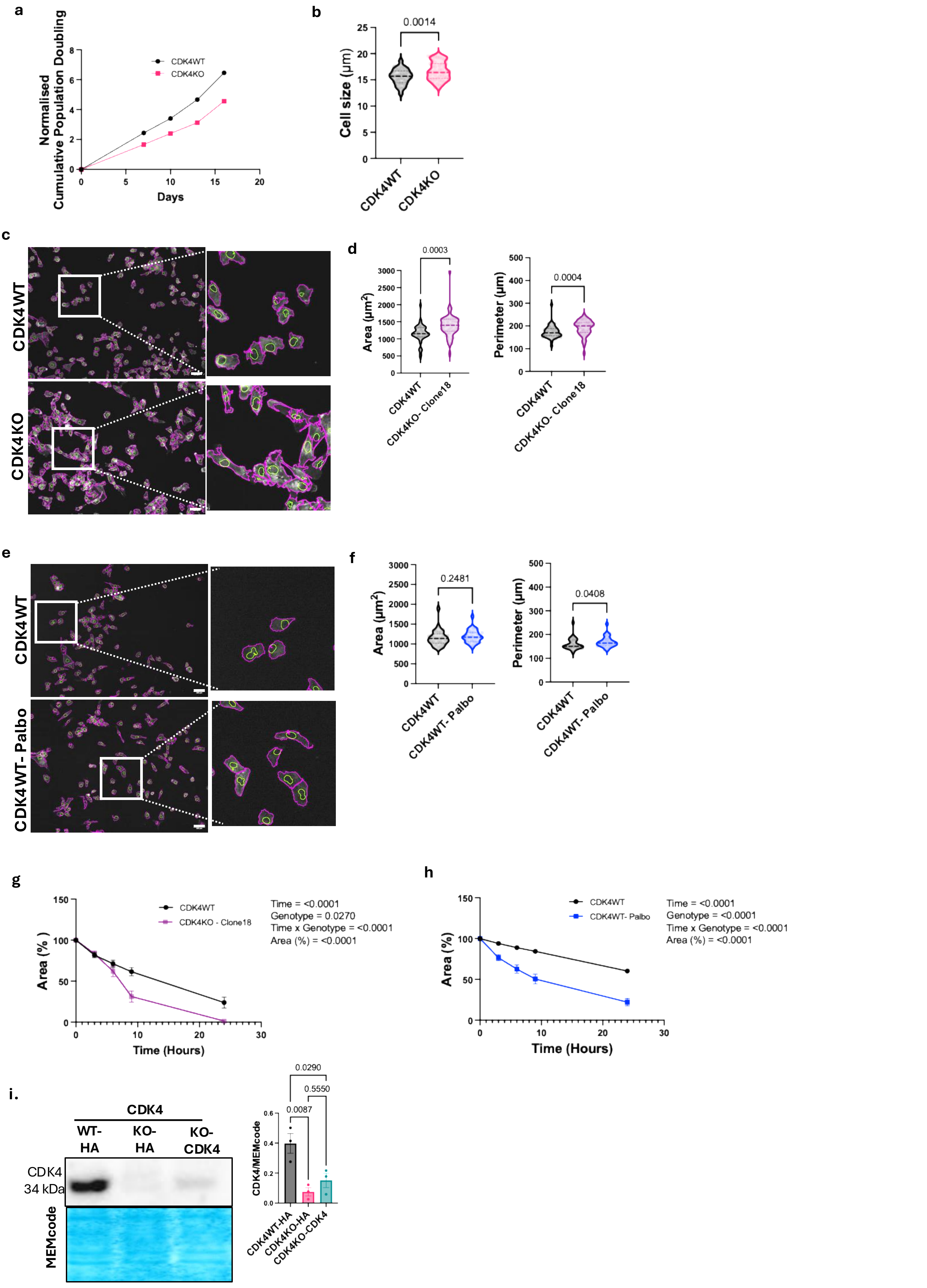
a
Normalised Cumulative Population Doubling
CDK4WT
CDK4KO
Days
b
0.0014
Cell size (µm)
CDK4WT
CDK4KO
c
CDK4WT
CDK4KO
d
0.0003
Area (µm²)
0.0004
Perimeter (µm)
CDK4WT
CDK4KO- Clone18
e
CDK4WT
CDK4WT- Palbo
f
0.2481
0.0408
Area (µm²)
Perimeter (µm)
CDK4WT
CDK4WT- Palbo
g
Area (%)
CDK4WT
CDK4KO - Clone18
Time = <0.0001
Genotype = 0.0270
Time x Genotype = <0.0001
Area (%) = <0.0001
Time (Hours)
h
Area (%)
CDK4WT
CDK4WT- Palbo
Time = <0.0001
Genotype = <0.0001
Time x Genotype = <0.0001
Area (%) = <0.0001
Time (Hours)
i.
CDK4
WT-HA
KO-HA
KO-CDK4
CDK4
34 kDa
MEMcode
0.0290
0.5550
0.0087
CDK4/MEMcode
CDK4WT-HA
CDK4KO-HA
CDK4KO-CDK4

**Sup. Figure 1. CDK4 inhibition or knockout (KO) increases cell size, induces topographic changes and promotes migration of TNBC cells.**

a. Representative graph for cumulative population doubling rate of CDK4WT and CDK4KO MDA-MB-231 TNBC cells from live cell counting during cell passaging for 20 days. b. Quantification of cell size by Trypan blue from live cell counting during cell passaging. c. Representative images (Sharp contrast) from high throughput celena microscope showing cell's boundaries in pink line for CDK4WT and another CDK4KO- clone 18 MDA-MB-231 cells, white square and dotted line showing the higher magnification. d. Associated quantification of morphological parameters (Celena Microscope): area and perimeter of CDK4WT and CDK4KO- clone 18 cells. N= 3 independent biological replicates. Unpaired t-test, Wilcoxon test. e-f. Representative images (Sharp contrast) and quantification of morphological parameters (Celena): area and perimeter of CDK4WT and CDK4WT- Palbociclib (Palbo, 1μM) treated cells, white square and dotted line showing the higher magnification. N= 3 independent biological replicates. Unpaired t-test, Man- Whitney test. g. Scratch Assay analysis (Area%) vs Time (hours) for CDK4WT and CDK4KO- clone 18. N= 3 independent biological replicates. 2-way ANOVA, Tukey's multiple comparison test. h. Scratch Assay analysis (Area%) vs Time (hours) for CDK4WT and CDK4WT- Palbo. N= 3 independent biological replicates. 2-way ANOVA, Tukey's multiple comparison test. i. Immunoblot showing the transfection of HA (empty plasmid vector) and CDK4-plasmid vector with the CDK4 expression and MEMcode, with associated quantification. One-way ANOVA, Tukey's multiple comparison test. Exact p-values are displayed.

**a**

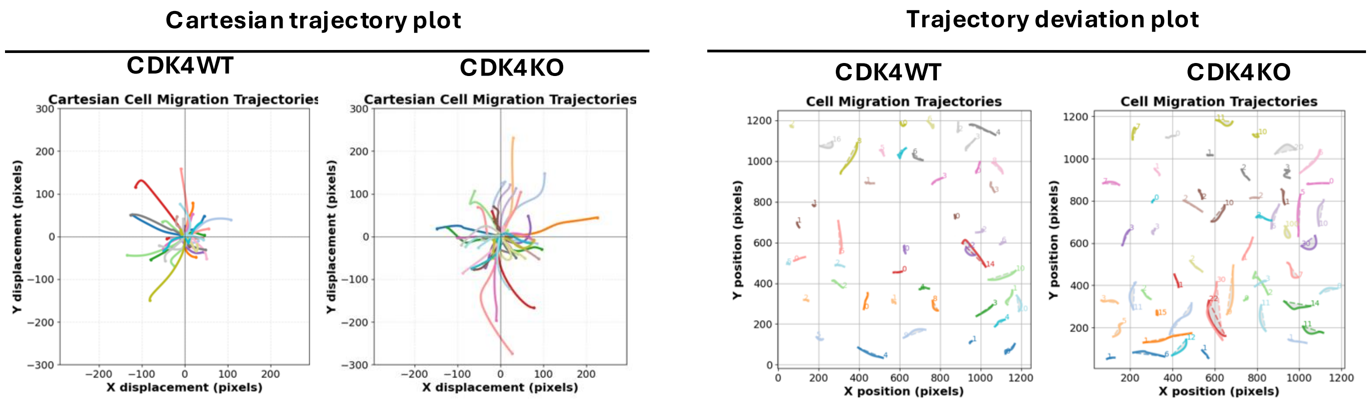
Cartesian trajectory plot
CDK4WT
CDK4KO
Cartesian Cell Migration Trajectories
X displacement (pixels)
Y displacement (pixels)
Trajectory deviation plot
Cell Migration Trajectories
X position (pixels)
Y position (pixels)

**b**

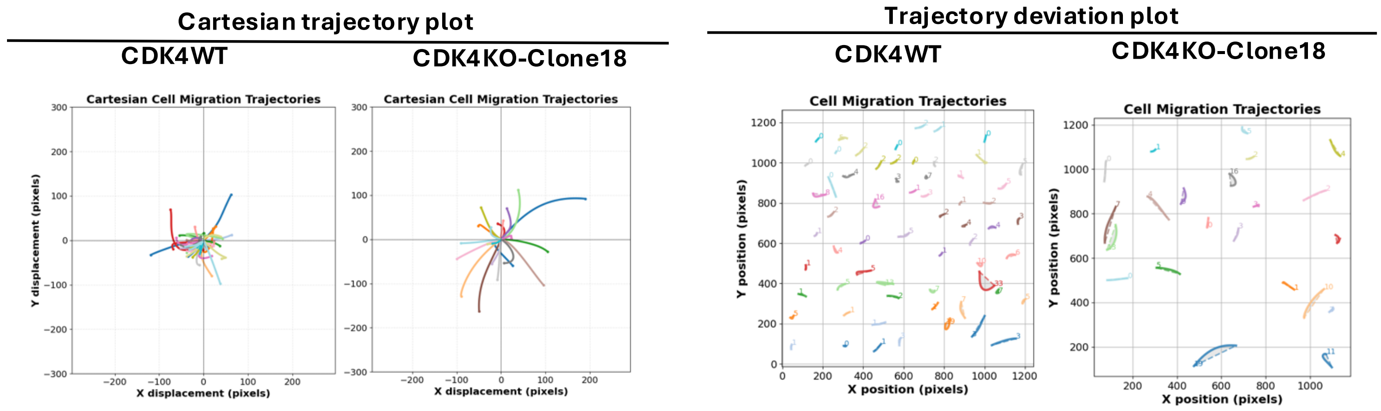
Cartesian trajectory plot
CDK4WT
CDK4KO-Clone18
Cartesian Cell Migration Trajectories
X displacement (pixels)
Y displacement (pixels)
Trajectory deviation plot
Cell Migration Trajectories
X position (pixels)
Y position (pixels)

**c**

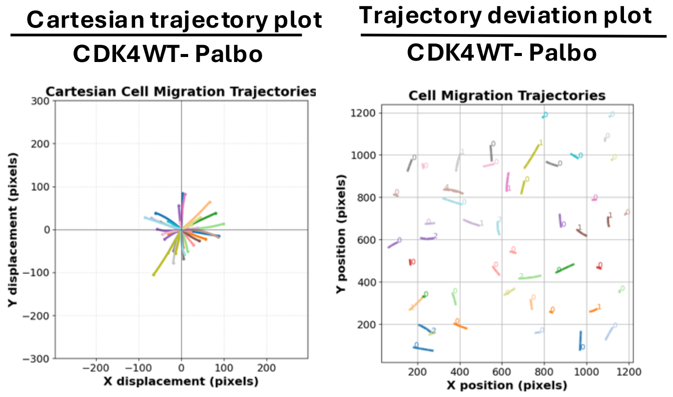
Cartesian trajectory plot
CDK4WT- Palbo
Cartesian Cell Migration Trajectories
X displacement (pixels)
Y displacement (pixels)
Trajectory deviation plot
CDK4WT- Palbo
Cell Migration Trajectories
X position (pixels)
Y position (pixels)

**d.**

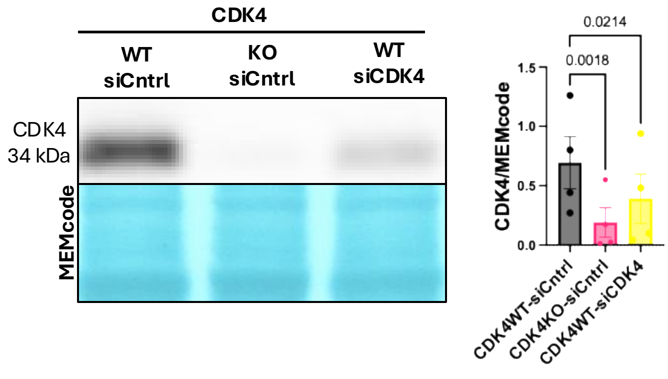
CDK4
WT siCntrl
KO siCntrl
WT siCDK4
CDK4 34 kDa
MEMcode
CDK4/MEMcode
0.0214
0.0018
CDK4WT-siCntrl
CDK4KO-siCntrl
CDK4WT-siCDK4

**e**

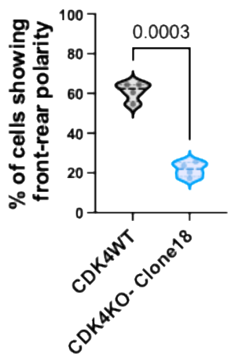
% of cells showing front-rear polarity
0.0003
CDK4WT
CDK4KO- Clone18

**Sup. Figure 2. CDK4KO disrupts the directional migration and front-rear polarity of actin localization in TNBC cells.**

a-b-c. Cartesian Trajectory visualization across experimental conditions, showing representative paths of multiple cells. Each line corresponds to the motion path of a single cell, aligned at the origin to allow comparison of migration patterns. Trajectory Deviation from a straight migration path. The original cell trajectory (solid line) is plotted alongside the straight-line displacement from the initial to the final position (dashed line). The area between these two paths (in light Gray) quantifies the deviation from directional movement, and its value is reported. For CDK4WT, CDK4KO- Clone 18 and CDK4WT- Palbo treated cells. d. Immunoblot showing the transfection of siCntrl and siCDK4 with the CDK4 expression and MEMcode, with associated quantification. One-way ANOVA, Tukey’s multiple comparison test. e. Quantification of % of cells showing front-rear polarity for CDK4WT and CDK4KO- Clone 18, N= 4 independent biological replicates, n= 50-60 cells. Paired t test. Exact p-values are displayed

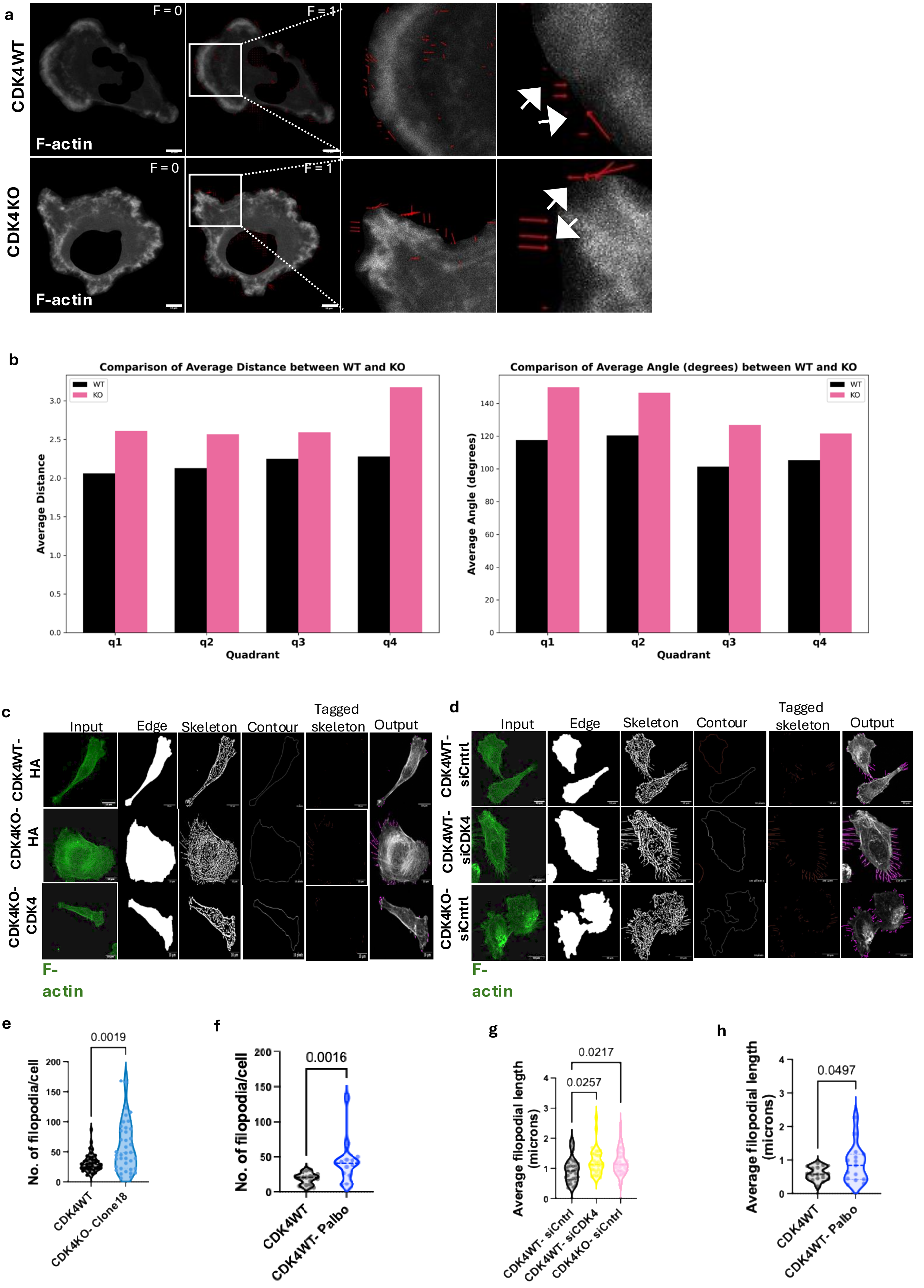
a
CDK4WT
CDK4KO
F = 0
F = 1
F-actin
b
Comparison of Average Distance between WT and KO
Average Distance
Quadrant
WT
KO
q1
q2
q3
q4
Comparison of Average Angle (degrees) between WT and KO
Average Angle (degrees)
c
Input
Edge
Skeleton
Contour
Tagged skeleton
Output
CDK4WT-HA
CDK4KO-HA
CDK4KO-CDK4
F-actin
d
CDK4WT-siCntrl
CDK4WT-siCDK4
CDK4KO-siCntrl
e
0.0019
No. of filopodia/cell
CDK4WT
CDK4KO- Clone18
f
0.0016
CDK4WT- Palbo
g
0.0217
0.0257
Average filopodial length (microns)
CDK4WT- siCntrl
CDK4WT- siCDK4
CDK4KO- siCntrl
h
0.0497

**Sup. Figure 3. CDK4 inactivation increases the expression of actin-associated proteins, altering actin dynamics and promoting filopodium formation in TNBC cells.**

a. Representative snapshots from motion vector analysis videos depict retraction (red arrows, inward) and expansion (white arrows, outward), with a 10 μm, scale bar. N= 3 independent biological replicates, n= 40-50 cells. b-c. Bar graphs showing the comparison of average distance and comparison of average angle (degree) of motion vector in each quadrant (q1, q2, q3, q4) for CDK4WT and CDK4KO. c. Image showing filopodial Quantification by using Filo Quant (plugin) from Fiji. Showing input image, edge, skeleton, contour, tagged skeleton, output for CDK4 transfection cell. d. Image showing filopodial Quantification by using Filo Quant from Fiji. Showing input image, edge, skeleton, contour, tagged skeleton, output for siCDK4 transfected cell. e. Quantification of No. of Filopodia/cell in CDK4WT and CDK4KO- clone 18 cells. Unpaired T-test Mann-Whitney test, n = approx. 30-40cells, N=3 independent biological replicates. f. Quantification of No. of Filopodia/cell in CDK4WT and CDK4WT- Palbociclib treated cells. Unpaired T-test Mann-Whitney test, n = approx. 30-40 cells, N=3 independent biological replicates. g. Quantification of Average length of Filopodia (microns) for siCDK4 transfected cell. n = approx. 30-40cells, N= 3 independent biological replicates. Kruskal-Wallis test, Dunn's multiple comparisons test. h. Quantification of Average length of Filopodia (microns) for CDK4WT and CDK4WT- Palbociclib treated cell, Unpaired t test, n = approx. 30-40 cells, N=3 independent biological replicates. Exact p-values are displayed.

**a**

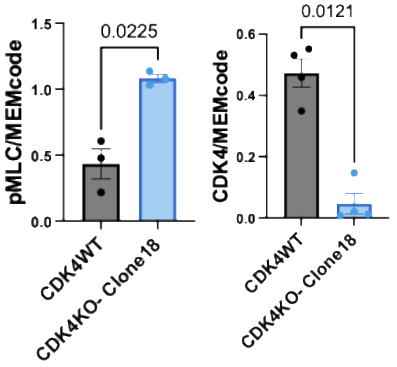


**b**

| Kinase | Kinase Group | Score Rank | Site Percentile | Percentile Rank | Gene Name |
|---|---|---|---|---|---|
| CDK4 | CMGC | 4 | **96.498459** | 4 | **MYO9B** |
| CDK4 | CMGC | 47 | 80.416994 | 65 | ARHGERF18 |
| CDK4 | CMGC | 22 | 92.113374 | 24 | ANLN |
| CDK4 | CMGC | 29 | 92.576298 | 43 | SPEN |
| CDK4 | CMGC | 45 | 94.923551 | 80 | EMD |
| CDK4 | CMGC | 274 | 18.786487 | 267 | CDC42EP1 |
| CDK4 | CMGC | 7 | 93.587962 | 9 | PKN1 |
| CDK4 | CMGC | 29 | 98.28972 | 29 | SH3BP1 |
| CDK4 | CMGC | 11 | 99.249411 | 9 | SYDE |

**c**

| Experiment | Target | Kinase | Kinase Group | Site Percentile | Percentile Rank | Site Sequence | Site |
|---|---|---|---|---|---|---|---|
| Phospho proteomics | Myo9b | CDK4 | CMGC | 96.498 | 4 | LKLGFSSPYEGVLNKS*PKTRDIQEEELEVLL | 1935 |
| Phospho proteomics | Myo9b | CDK4 | CMGC | 84.885 | 24 | LKLGFSS*PYEGVLNKSPKTRDIQEEELEVLL | 1926 |

**d**

```
sp|Q13459|MYO9B_HUMAN               VKMEEISQLEAAESIAFRRLSLLRQNAPWPLKLGFSSPYEGVLNKSPKTRDIQE--EELE    1947
tr|H9FUR6|H9FUR6_MACMU              VKMEEISQLEAAESIAFRRLSLLRQNALWPLKLGFSSPYEGVLNKSPKARDSQG--EELE    1947
sp|Q9QY06|MYO9B_MOUSE               MKMEEINHLEAAESIAFRRLSLLRQNAPWPLKLGFSSPYEGVRIKSPRTPVVQD--L-EL    1904
sp|Q63358|MYO9B_RAT                 VKMEEINHLEAAESIAFRRLSLLRQNAPWPLKLGFSSPYEGVRTKSPRTPVVQD--LEEL    1907
tr|A0A1V4KQM6|A0A1V4KQM6_PATFA      VKMDEINQLEAAESIAFRRLSLLRQNTLWPVKLGFSSPYEGMLSKSSQAKGGDSGSSELD    1967
tr|A0A8V1AFP8|A0A8V1AFP8_CHICK      IKMDEINQLEAAESIAFRRLSLLRQNTLWPVKLGFSSPYEGMLSKSPQVKGNDSGSSELD    1994
tr|K7F272|K7F272_PELSI              LKMDEINQLEVAESIAFRRLSLLRQNMLWPVKLGFSSPFEGVLCKSPQDKGNDNNFPKLE    1957
tr|A0A6I8QUN6|A0A6I8QUN6_XENTR      IKMQEIGLLEAAEIRAVTRLSLLRNTTSKSQ-----------------ASNGKSLPDLD    1918
tr|A0A8M3AQC3|A0A8M3AQC3_DANRE      QKMEEIQQLESAEAMAVKELQLRRQNTIFEPPKS---------------DSNASS-----    1824
                                     **:**  ** **  *. .*.* *:.
```

* Fully conserved residue (exact amino acid in all sequences at that position)
: Strong conservation (residues are different but have strongly similar properties)
. Weak conservation (residues have weakly similar properties)

**Sup. Figure 4. CDK4KO enhances RhoA activity and ROCK-dependent migration, with Myo9b phosphorylation as a CDK4 target**

a. Immunoblot Quantification for CDK4WT and CDK4KO- clone 18 cells for pMLC, CDK4, normalize to MEMcode. Paired t test, N= 4 independent biological replicates, Exact p-values are displayed. b-c. Excel sheet -image for kinase prediction from Phosphosite. Org. database d. Sequence alignment by Align-UniProt for Myo9b across species, highlighted for S1926 (red) and S1935 (pink).

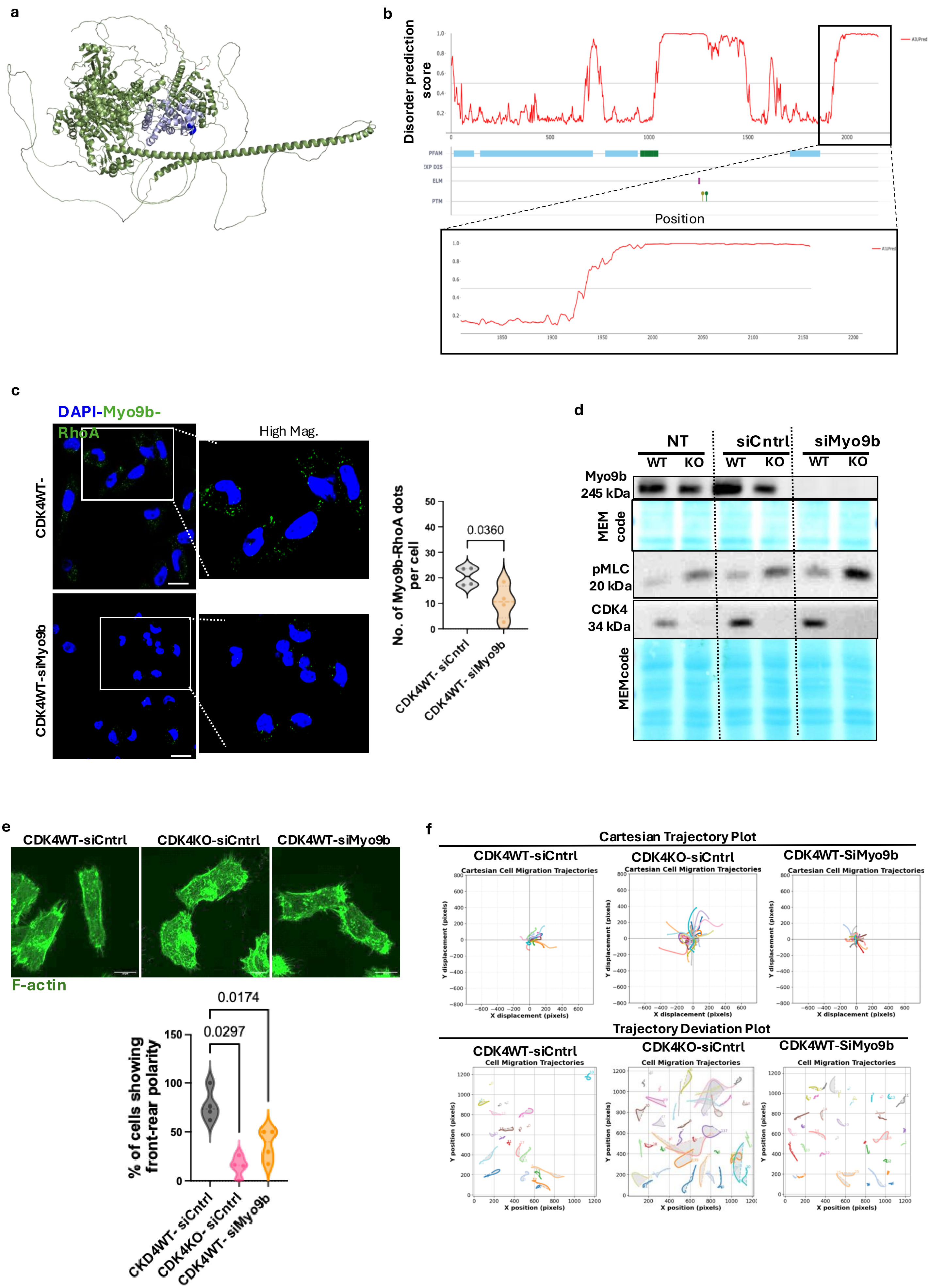

a
b
Disorder prediction score
Position
PFAM
ELM
PTM
AIUPred
c
DAPI-Myo9b-RhoA
High Mag.
CDK4WT-
CDK4WT-siMyo9b
No. of Myo9b-RhoA dots per cell
0.0360
CDK4WT- siCntrl
CDK4WT- siMyo9b
d
NT
siCntrl
siMyo9b
WT
KO
Myo9b 245 kDa
MEM code
pMLC 20 kDa
CDK4 34 kDa
MEMcode
e
CDK4WT-siCntrl
CDK4KO-siCntrl
CDK4WT-siMyo9b
F-actin
% of cells showing front-rear polarity
0.0174
0.0297
CKD4WT- siCntrl
CDK4KO- siCntrl
CDK4WT- siMyo9b
f
Cartesian Trajectory Plot
CDK4WT-siCntrl
CDK4KO-siCntrl
CDK4WT-SiMyo9b
Cartesian Cell Migration Trajectories
Y displacement (pixels)
X displacement (pixels)
Trajectory Deviation Plot
Cell Migration Trajectories
Y position (pixels)
X position (pixels)

**Sup. Figure 5. Phosphorylated Myo9b at S1935 regulates RhoA activity via the interaction between Myo9b-RhoGAP and RhoA**

a. Human Myo9b (Q13459 in UniProt) structural prediction by AlphaFold. The RhoGAP domain is shown in purple, and within the RhoGAP, the RhoA interaction site reported in UniProt is shown in dark blue. b. AIUPred prediction of disordered regions for Myo9b, with a zoomed-in view (bottom) of the terminal disordered region. c. Proximity Ligation Assay (PLA) using Myo9b and RhoA antibodies in CDK4WT-siCntrl and CDK4WT-siCntrl TNBC cells. Representative pictures and associated quantification of Myo9-RhoA dots per cell. Scale bar: 10 [illegible]m. N = 4 independent biological replicates, n=50-60 cells. Two-sided paired T-test. d. Qualitative observation by immunoblot for CDK4WT and CDK4KO, siCntrl and siMyo9b transfected cells for Myo9b, pMLC, and CDK4 check. Blot for Myo9b (6%) and pMLC, CDK4 (12%) were done on separate gels. e. Representative images (Sharp contrast) and quantification of % of cells showing front-rear polarity in sicntrl and siMyo9b treated CDK4WT and CDK4KO MDA-MB-231 cells. N= 3 independent biological replicates, n= 30-40 cells, approx. One-way ANOVA, Tukey's multiple comparison test. f. Cartesian Trajectory visualization across experimental conditions, showing representative paths of multiple cells. Each line corresponds to the motion path of a single cell, aligned at the origin to allow comparison of migration patterns. Trajectory Deviation from a straight migration path. The original cell trajectory (solid line) is plotted alongside the straight-line displacement from the initial to the final position (dashed line). The area between these two paths (in light Gray) quantifies the deviation from directional movement, and its value is reported. For CDK4WT, CDK4KO- siCntrl and siMyo9b transfected cells. Exact p-values are displayed.

**Table S1**. Residues separated by 10Å or less between Myo9b and RhoA in their different phosphorylated forms.

| Structure | Chain A residue | Chain B residue | Distance (Å) |
|---|---|---|---|
| Myo9b_P1935-RhoA | 1739 | 90 | 4.592984 |
| Myo9b_P1935-RhoA | 1736 | 14 | 4.8282957 |
| Myo9b_P1935-RhoA | 1735 | 14 | 5.2011657 |
| Myo9b_P1935-RhoA | 1738 | 90 | 5.5369062 |
| Myo9b_P1935-RhoA | 1736 | 15 | 5.55995 |
| Myo9b_P1935-RhoA | 1845 | 35 | 5.754721 |
| Myo9b_P1935-RhoA | 1763 | 134 | 5.772862 |
| Myo9b_P1935-RhoA | 1735 | 15 | 5.8491893 |
| Myo9b_P1935-RhoA | 1737 | 14 | 5.990028 |
| Myo9b_P1935-RhoA | 1845 | 36 | 5.998946 |
| Myo9b_P1926-RhoA | 1594 | 73 | 5.4030566 |
| Myo9b_P1926-RhoA | 1586 | 40 | 5.620079 |
| Myo9b_P1926-RhoA | 1590 | 41 | 5.7992363 |
| Myo9b_P1926-RhoA | 1587 | 38 | 5.853452 |
| Myo9b_P1926-RhoA | 1591 | 73 | 6.0960913 |
| Myo9b_P1926-RhoA | 1586 | 38 | 6.1670833 |
| Myo9b_P1926-RhoA | 1598 | 75 | 6.2774515 |
| Myo9b_P1926-RhoA | 1590 | 40 | 6.382666 |
| Myo9b_P1926-RhoA | 1530 | 65 | 6.4272532 |
| Myo9b_P1926-RhoA | 1591 | 72 | 6.4598956 |
| Myo9b_P1926P1935-RhoA | 1478 | 169 | 4.822135 |
| Myo9b_P1926P1935-RhoA | 1496 | 42 | 5.1538534 |
| Myo9b_P1926P1935-RhoA | 1489 | 46 | 5.193765 |
| Myo9b_P1926P1935-RhoA | 1436 | 118 | 5.3936844 |
| Myo9b_P1926P1935-RhoA | 1594 | 73 | 5.4277043 |
| Myo9b_P1926P1935-RhoA | 1437 | 120 | 5.5397587 |
| Myo9b_P1926P1935-RhoA | 1485 | 46 | 5.5708995 |
| Myo9b_P1926P1935-RhoA | 1494 | 44 | 5.5948105 |

| | | | |
|---|---|---|---|
| Myo9b_P1926P1935-RhoA | 1586 | 40 | 5.6012635 |
| Myo9b_P1926P1935-RhoA | 1490 | 26 | 5.608255 |
| Myo9b-RhoA | 1493 | 44 | 4.335871 |
| Myo9b-RhoA | 1489 | 47 | 4.4259806 |
| Myo9b-RhoA | 1496 | 42 | 4.6845984 |
| Myo9b-RhoA | 1491 | 46 | 4.7287345 |
| Myo9b-RhoA | 1488 | 48 | 4.8025928 |
| Myo9b-RhoA | 1492 | 45 | 5.07927 |
| Myo9b-RhoA | 1491 | 45 | 5.520493 |
| Myo9b-RhoA | 1487 | 49 | 5.538419 |
| Myo9b-RhoA | 1488 | 47 | 5.5744896 |
| Myo9b-RhoA | 1594 | 73 | 5.575784 |

### **Table S2.** List of Antibodies

| Antibodies | Dilution (WB) | Dilution (IF) | Dilution (PLA) | Species | Reference |
|---|---|---|---|---|---|
| Actin (Act) | 1:1000 | | | Rabbit | Sigma- Aldrich (A2066) |
| Tubulin (Tub) | 1:1000 | 1:1000 | | Mouse | Sigma- Aldrich (T6199) |
| RhoA (RHOA Monoclonal Antibody 1B12) | 1:1000 | | 1:1000 | Mouse | Thermofisher Life Technoligies # H00000387-M04 |
| Phospho Myosin Light Chain2 (Ser19) (pMLC) | 1:1000 | | | Rabbit | Cell Signaling Technology (CST) #3671 |
| Myosin Light Chain 2 (D18E2) (Total MLC) | 1:1000 | | | Rabbit | Cell Signaling Technology (CST) #8505 |
| Phospho Retinoblastoma (pRb) (s780) | 1:1000 | | | Rabbit | Cell Signaling Technology (CST) #8180 (D59B7) |
| Retinoblastoma (Total Rb) | 1:1000 | | | | Scanta Cruz (SC-50) (C-15) |
| Cyclin Dependent Kinase 4 (CDK4) | 1:1000 | | | Rabbit | Cell Signaling Technology (CST) #1790 (D9G3E) |
| Myo9b (Myosin IX b) | 1:1000 | | 1:1000 | Rabbit | Proteintech (LubioScience GmbH) Cat n° 12432-1-AP |
| Actin- Phalloidin (green) | 1:500 | | | | ThermoFisher Scientific (A12379) Alexa Fluor ™ 488) |
| Hoescht 33342 | | 1:10000 | | | (H21492) Hoechst 33342, Trihydrochloride, Trihydrate - FluoroPure™ Grade |
